# GPC: An expressive and tractable deep generative model for genetic variation data

**DOI:** 10.1101/2023.05.16.541036

**Authors:** Prateek Anand, Anji Liu, Meihua Dang, Boyang Fu, Xinzhu Wei, Guy Van den Broeck, Sriram Sankararaman

## Abstract

Generative models play an important role in population genetics, being used to generate artificial genomes (AGs) that help benchmark methods, test evolutionary hypotheses, and construct reference panels for imputation while working around data-sharing restrictions. Existing generative models of genetic variation, however, struggle to faithfully express dependencies in the data while retaining tractability and preserving privacy.

We introduce Genetic Probabilistic Circuits (GPC), a deep generative model for genetic variation data based on hidden Chow-Liu trees represented as probabilistic circuits. GPC generalizes traditional hidden Markov models by allowing arbitrary tree structures over latent variables, enabling it to capture long-range dependencies among SNPs. GPC is tractable, supporting exact computation of marginal and conditional probabilities that enables both AG generation and direct genotype imputation (avoiding the need for simulating AGs).

We show that GPC generates AGs that accurately reproduce population structure and linkage disequilibrium patterns across a range of length scales. Compared to other deep generative approaches, GPC consistently improves imputation accuracy, with particularly strong gains for low-frequency variants and in populations that are not well-represented in public reference panels. Finally, we show that GPC better preserves privacy of the training data, thereby providing a practical framework for AG generation in settings with limited data access.

## 1 Introduction

Generative models of genetic variation play a central role in population genomics [1, 2]. By modeling dependencies across individuals and variants, these models support core analyses such as genotype imputation [3], haplotype phasing [4], and ancestry inference [5]. They also form the foundation of simulators that generate artificial genomes (AGs) [6, 7, 8, 9, 10], which are widely used to test evolutionary hypotheses, estimate population genetic parameters, validate empirical results, and benchmark new methods. As the sharing of primary genetic data becomes increasingly restricted due to privacy and consent constraints, accurate simulation tools have become essential for reproducibility and equity. Trained generative models, or data simulated from them, can in principle be shared without requiring access to identifiable genomic data.

Classical population genetic simulators rely on the coalescent model [11], which generates genomes based on demographic history together with mutation and recombination rates. The coalescent with recombination [12] connects observed genetic variation to latent genealogies that change along the genome [13] and is highly expressive, but exact inference under this model is computationally challenging, motivating tractable approximations. One class approximates the coalescent as a Markovian process along the genome [14, 15], forming the basis of widely used simulators such as `msprime` [9] and its extensions [10]. A second class dispenses with explicit genealogical modeling and instead directly approximates the data distribution. A prominent example is the product-of-approximate-conditionals (PAC) model [16], which naturally yields a hidden Markov model (HMM) [17]. HMM-based models have proven highly successful in applications including haplotype phasing [18, 19], genotype imputation [20, 3], and ancestry inference [21, 22].

Motivated by the success of deep learning, recent work has explored deep generative models as an alternative approach to represent genetic variation [23, 24, 25]. Models based on generative adversarial networks (GANs) [25, 26, 27, 28], variational autoencoders (VAEs) [23, 29, 30], restricted Boltzmann machines (RBMs) [25], and more recently diffusion models [31] can be more expressive than HMMs and often produce AGs that visually resemble real genomes in principal component analysis (PCA) or allele frequency summaries. However, these models have critical limitations for genomic applications. First, GANs do not define a probability distribution over the data, making likelihood-based inference impossible. RBMs define a distribution but require computing an intractable partition function over exponentially many configurations, preventing exact likelihood evaluation. VAEs define a probability distribution but the marginal likelihood requires integrating over a continuous latent space with a nonlinear decoder, which is intractable; only a lower bound (the ELBO) is accessible in practice. Diffusion models can in principle compute exact likelihoods via probability flow ordinary differential equations, but scale poorly to SNP data: existing approaches [31] require lossy dimensionality reduction (e.g., per-gene PCA) as a preprocessing step before the model operates [31], and have not been evaluated on imputation tasks. Second, none of these approaches support efficient conditional probability estimation. This limitation makes model comparison difficult and limits their use in downstream tasks such as genotype imputation, which require computing the probability of unobserved variants conditioned on observed variants. A common workaround involves generating AGs from these models, which then serve as a reference panel for imputation methods such as Impute5, Beagle, or Minimac [32, 33, 34], though this approach can introduce additional noise in the imputation process. Third, training these models is challenging due to complex optimization objectives and large hyperparameter spaces, and convergence must be assessed through subjective visual inspection rather than principled likelihood monitoring. These difficulties collectively highlight the need for deep generative models that are both expressive and tractable.

We address this need by introducing GPC (Genetic Probabilistic Circuits), a class of deep generative models that fits genetic variation data with high accuracy while supporting efficient and exact inference. To model the distribution over single nucleotide polymorphisms (SNPs) in an individual, we propose a latent variable model in which each SNP is associated with a hidden variable and the hidden variables form a tree-structured graphical model. This model, termed the hidden Chow-Liu tree (HCLT) [35], generalizes the HMM. Classical HMMs impose a chain structure where hidden variables corresponding to consecutive SNPs are adjacent in the tree. HCLTs relax this constraint by allowing arbitrary tree structures, which enables the model to capture long-range dependencies, or linkage disequilibrium (LD), across the genome [36]. This flexible structure allows HCLTs to more faithfully represent the distribution of genetic variation. While HCLTs are more expressive than HMMs, naively training and performing inference in HCLTs at the scale required for modern genomic datasets would be computationally prohibitive. Our approach represents HCLTs as probabilistic circuits (PCs) [37, 38], a circuit-based framework that permits tractable inference tasks such as exact likelihood and conditional probability computation. This representation allows us to leverage stochastic gradient-based learning and GPU acceleration to train HCLTs with millions of parameters on large genomic datasets.

GPC, like GANs and RBMs, can generate AGs for use as reference panels with standard imputation tools. Unlike these methods, GPC can also perform imputation directly through efficient computation of conditional probabilities, without the need for generating AGs as an intermediate step. We show that this ability further improves imputation accuracy.

We first compare GPC to baselines on genomic regions chosen from the 1000 Genomes Project (1KG) and UK Biobank (UKBB), demonstrating that GPC achieves higher held-out log-likelihoods than simpler probabilistic models and generates AGs that accurately capture population structure and linkage disequilibrium patterns across all distance ranges. We then evaluate GPC extensively for genotype imputation. Across both datasets, GPC yields significant improvements in imputation accuracy compared to other deep generative models, with substantial improvements in settings where reference genomes from the target population are not publicly available. In such population-specific settings, GPC consistently outperforms other deep generative methods, with particularly strong gains for low-frequency variants. Finally, we find that AGs produced by GPC offer improved privacy properties compared to previous deep generative models. Overall, our results indicate that the increased expressivity and tractability of GPC lead to more accurate and privacy-preserving models of human genetic variation.

## 2 Results

### 2.1 Methods Overview

We present GPC, a deep generative model for genetic variation data. GPC is based on hidden Chow-Liu trees (HCLTs) [35], a class of latent variable models that generalize hidden Markov models (HMMs), represented as probabilistic circuits (PCs) [37, 38] for efficient inference and learning.

#### Hidden Chow-Liu Trees

HCLTs are tree-structured graphical models that represent the probability distribution over observed random variables, **X** = (*X*_1_,…, *X*_*N*_). In the context of genetic variation across *N* SNPs, each observed *X*_*n*_ represents the genotype at SNP *n* (e.g., *X*_*n*_ ∈ {0, 1} for haploid genomes). HCLTs rely on the following key designs:

- Hidden Variable Mapping: For every observed variable *X*_*n*_, the model introduces a corresponding hidden variable *Z*_*n*_ (which takes one of *L* discrete values). Together, these form hidden variables **Z** = (*Z*_1_,…, *Z*_*N*_).
- Chow-Liu Tree Architecture: Each observed variable *X*_*n*_ connects solely to its hidden counterpart *Z*_*n*_. The hidden variables, however, connect to each other to form a tree structure. This tree is learned using the Chow-Liu algorithm [39] to capture the strongest pairwise correlations between variables (Figure 1).
- Parameterization: The entire HCLT is defined by just two sets of conditional probability distributions: the emissions *P* (*X*_*n*_|*Z*_*n*_) and the tree transitions *P* (*Z*_*n*_|*Z*_Pa(*n*)_), where Pa(*n*) denotes the parent of node *n* in the hidden variable tree.

**Figure 1:**
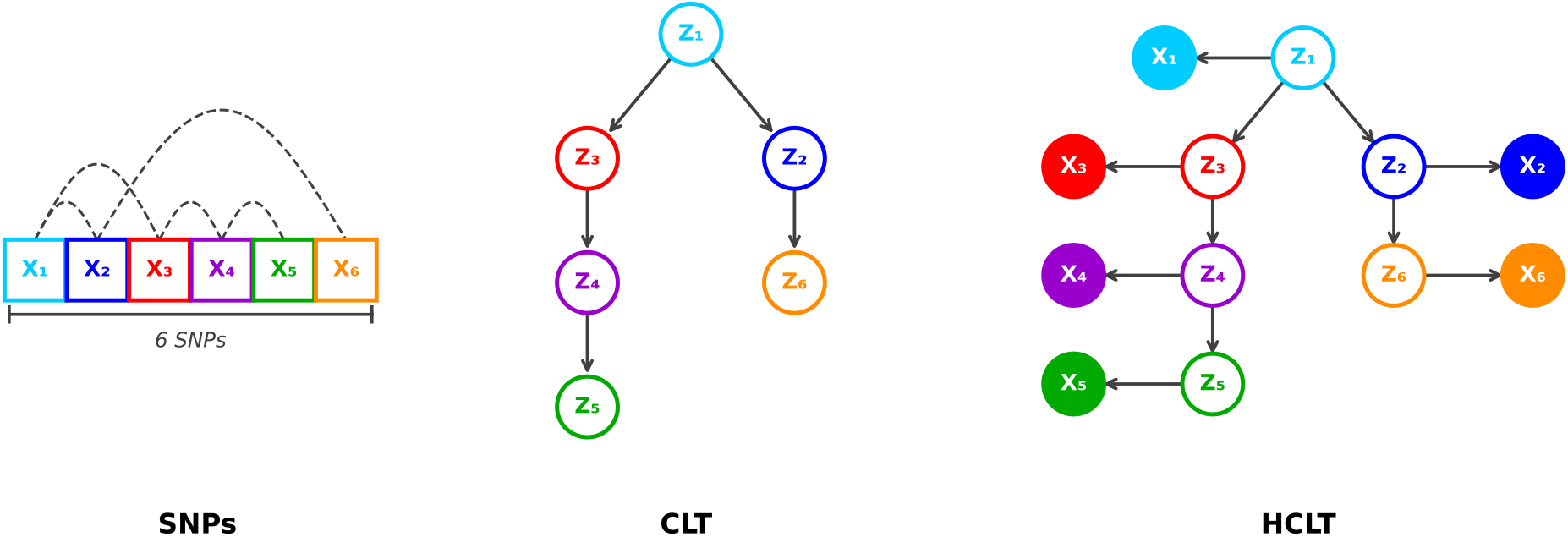
Generating HCLT structures given genetic data. **SNPs:** An example dataset of 6 SNPs (*X*_1_,…, *X*_6_), where dashed lines indicate SNPs with strong correlations. **CLT:** A Chow–Liu Tree learned from the SNP data, represented as a Bayesian network over latent variables *Z*_1_,…, *Z*_6_. The tree structure places latent variables with stronger dependencies closer together, capturing the dominant correlation structure among the SNPs. **HCLT:** The Hidden CLT constructed from the CLT, in which each latent variable *Z*_*i*_ is additionally connected to its corresponding observed SNP *X*_*i*_ as a child node. The resulting structure respects the dependencies among SNPs while providing a principled generative model over the observed data.

The critical distinction from HMMs is structural. Classical HMMs impose a chain structure where hidden variables corresponding to consecutive SNPs are adjacent in the tree (*Z*_1_ → *Z*_2_ → … → *Z*_*N*_). This chain topology can only propagate information between distant SNPs through all intermediate variables. HCLTs relax this constraint by allowing arbitrary tree structures, so that SNPs with strong long-range correlations (such as *X*_1_ and *X*_5_ being highly correlated without being correlated with intermediate SNPs) can be placed close together in the tree (Figure 1). This flexible structure allows HCLTs to more faithfully capture linkage disequilibrium patterns at all length scales.

#### Probabilistic Circuits and Tractable Inference

While HCLTs are more expressive than HMMs, naively training and performing inference in HCLTs at the scale required for modern genomic datasets would be computationally prohibitive. Our key insight is to represent HCLTs as probabilistic circuits (PCs) [37, 38]. A PC represents a joint probability distribution through a directed acyclic graph (DAG) consisting of input, sum, and product nodes. Product nodes define factorized distributions over their inputs, while sum nodes represent mixtures weighted by learnable parameters. Under structural constraints known as smoothness and decomposability, arbitrary marginal queries can be computed in time linear in the size of the circuit. This tractability is what enables GPC to perform three key operations efficiently: (1) computing likelihoods on held-out data for principled model comparison, (2) generating artificial genomes via ancestral sampling, and (3) performing direct genotype imputation through exact conditional probability computation. By representing HCLTs as equivalent PCs and training them using the PC package PyJuice [40], we can extensively parallelize Expectation-Maximization (EM) updates on GPUs, enabling efficient training even for models with over 88 million parameters (Section 4.2).

Generating AGs from GPC is performed via ancestral sampling through the circuit in time linear in the circuit size. For genotype imputation, GPC computes conditional probabilities *p*(*X*_missing_|*X*_observed_) directly as ratios of marginal queries, both of which can be evaluated via a single feedforward pass through the circuit. This direct imputation capability, unique to GPC among the deep generative approaches we consider, eliminates the need for intermediate AG generation and, as we show, yields consistently higher accuracy (see Section 4.2.2 for full details on sampling and conditional inference).

### 2.2 Data

Our experiments evaluate different aspects of GPC in terms of its ability to generate realistic AGs, its ability to perform genotype imputation, and its privacy properties. We perform the evaluations on three datasets with distinct properties: 1000 Genomes Project Phase 3 (1KG) [41], UK Biobank (UKBB) [42], and high-coverage 1KG (high-coverage 1KG) [43], with the high-coverage 1KG dataset only being used for the array-based imputation experiment (see Section 4.1 for details).

### 2.3 Evaluation

#### Baselines

We compare GPC against two categories of baselines. The first category consists of deep generative models that have been previously proposed for AG generation: (1) generative adversarial networks (WGAN) and (2) restricted Boltzmann machines (RBM) as implemented in [25]. We focus our comparison on these two methods as they are the most established deep generative approaches specifically designed and benchmarked for the local genetic structure and imputation tasks we evaluate here. More recent approaches including VAEs [29, 30] and diffusion models [31] have been proposed for related but distinct tasks, and do not support efficient, tractable probabilistic inference; we leave systematic comparison with these methods to future work. For both WGAN and RBM baselines, we use the samples generated by the corresponding authors for comparison^1^ in our 1KG experiments, but we also retrain their models on our own training split to ensure a fair comparison. In imputation experiments, we additionally compare against Impute5 [32], a state-of-the-art imputation tool widely used in genetics. Impute5 serves a dual role: it is used with real reference genomes to establish performance benchmarks, and it is used with AG reference panels generated by each method to evaluate their utility for imputation.

The second category consists of simpler probabilistic graphical models (PGMs) that support tractable likelihood computation: fully-factorized distributions (Indep), Markov chain models of order 1 (Markov), and non-homogeneous hidden Markov models (HMM). These serve as baselines for density estimation and AG quality comparisons.

#### Evaluation criteria

We evaluate these models using the following metrics: (1) log-likelihood on test data to assess the capability of each model as a density estimator; (2) summaries of AGs sampled from each model, including the top principal components and linkage disequilibrium at pairs of SNPs; (3) genotype imputation, which evaluates the models’ utility on an important downstream task; and (4) privacy analysis of the AGs sampled from each model.

### 2.4 GPC trains efficiently with objective convergence criteria

Training GPC requires minimal hyperparameter tuning. We set the number of latent states to 128 (the maximum feasible given memory constraints) and use a small pseudocount (0.005) for smoothing. Models were trained for 2,000–5,000 epochs depending on dataset size. We use the PC package PyJuice [40] to train GPC efficiently on GPUs, achieving EM updates in under 2, 10, and 3 seconds per epoch on the 1KG, UKBB, and high-coverage 1KG datasets, respectively. On a single NVIDIA RTX A5000 GPU with 24GB available memory, GPC trains in roughly 2, 6, and 4 hours on each dataset, respectively. Once the model is loaded, generating a single sample can be done in under 2, 5, and 3 seconds for each dataset, respectively. Imputing all missing SNPs within a single sample can be done in 20 milliseconds. A key advantage over other deep generative approaches is that convergence can be monitored via held-out log-likelihood, providing an objective stopping criterion rather than subjective visual inspection. Full training details for GPC and all baseline methods are provided in Section 4.4.

### 2.5 GPC accurately reconstructs local genetic structure

We first compared the ability of different generative models to represent genetic variation in the 1KG and UKBB datasets. We simulate AGs with GPC and all five baselines (WGAN, RBM, HMM, Indep, and Markov) for comparison. For both datasets, we use an 80% train / 20% test split performed at the individual level before haplotype separation; we generate 5,008 AGs from each model for 1KG and 10,000 AGs for UKBB.

GPC learns more accurate probabilistic models than HMM, Markov, and Indep as measured by their log likelihood on the test dataset that was not used for model fitting (Table 1). Note that we do not compare with WGAN and RBM since they do not support tractable exact likelihood computation. We additionally evaluated the quality of AGs based on whether they preserve distances across pairs of haploid genomes. To do this, we compute the pairwise differences of haploid genomes within a single dataset or between the test dataset and an AG dataset and compute the Wasserstein distance between these pairs of distributions; a lower Wasserstein distance indicates that the AGs tend to be more similar to real genomes in the test dataset. WGAN, RBM, and GPC all capture the distribution well with GPC having second-lowest distance behind WGAN on 1KG and having the lowest distance on UKBB (Figure S1 and rows 1-2 of Table S1).

**Table 1:** Log-likelihoods of probabilistic models that support tractable likelihood computation in 1KG and UKBB datasets. Averaged training and test log-likelihoods for Indep, Markov HMM, and GPC. The bold values highlight the best averaged log-likelihoods.

| Dataset | Category | INDEP | MARKOV | HMM | GPC |
| --- | --- | --- | --- | --- | --- |
| **1KG** | **train LL** | -2386.81 | -1806.33 | -591.08 | **-202.51** |
|  | **test LL** | -2404.51 | -1819.96 | -599.88 | **-265.06** |
| **UKBB** | **train LL** | -1642.62 | -1360.86 | -554.88 | **-120.10** |
|  | **test LL** | -1648.03 | -1362.16 | -554.38 | **-127.75** |

We then analyze the quality of AGs generated by all models in terms of capturing commonly-used summaries of genetic variation data. To visualize all methods in the same latent space, we merge eight datasets (training set, test set, and AGs from six methods) and apply a single PCA to the combined data. Inspecting the top six principal components (PCs) of genomes in the test set and the AGs, we find that AGs generated from GPC qualitatively capture the dominant structure in this dataset as do the deep learning methods (WGAN and RBM), unlike Indep, Markov, and HMM (Figure 2 for 1KG and Figure S2 for UKBB). To quantify the accuracy of these summaries, we compute the Wasserstein distance between the 2D PCA representations of the test genomes versus the AGs (Table S1) and find that distances for the deep learning methods (WGAN, RBM, and GPC) tend to be lower than simpler models.

**Figure 2:**
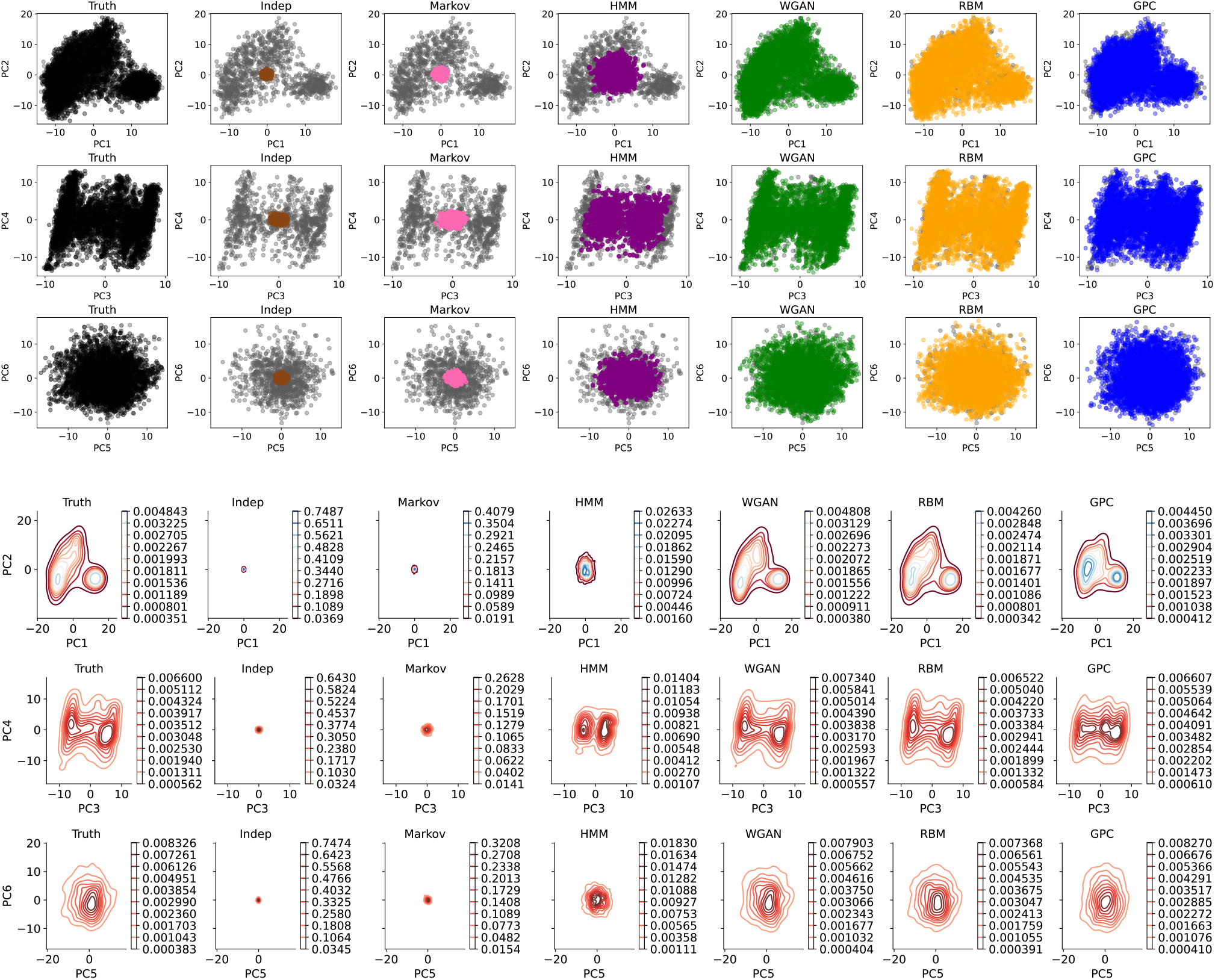
Principal components analysis for AGs generated by models trained on the 1KG dataset. (Top half) The top six principal components (PCs) of the held-out test set of genomes (gray) vs. AGs generated by each of Indep (brown), Markov (pink), HMM (purple), WGAN (green), RBM (orange), and GPC (blue). The leftmost plot shows the PCs of genomes from the training set as “perfectly” generated data (black). (Bottom half) A density map of the PCs for each dataset.

We also examined patterns of linkage disequilibrium (LD) to assess how the pairwise short and long-range correlations of SNPs can be captured by AGs. While WGAN and RBM tend to be accurate at longer length scales and HMM and Markov are accurate at shorter length scales, GPC is accurate across all length scales (Figure 3). Finally, we examined the topology of the CLT learned on the 1KG dataset to gain intuition into why GPC captures LD at all scales. In contrast to the chain-structured HMM in which every node has degree at most 2 and every edge connects adjacent SNPs along the genome, the learned CLT has 4,599 leaf nodes, 397 nodes with degree ≥ 5, and a maximum degree of 161, with 18.3% of edges connecting SNPs more than 1,000 positions apart in sequential index (Figure 4; Supplementary Figures S3 and S4).

**Figure 3:**
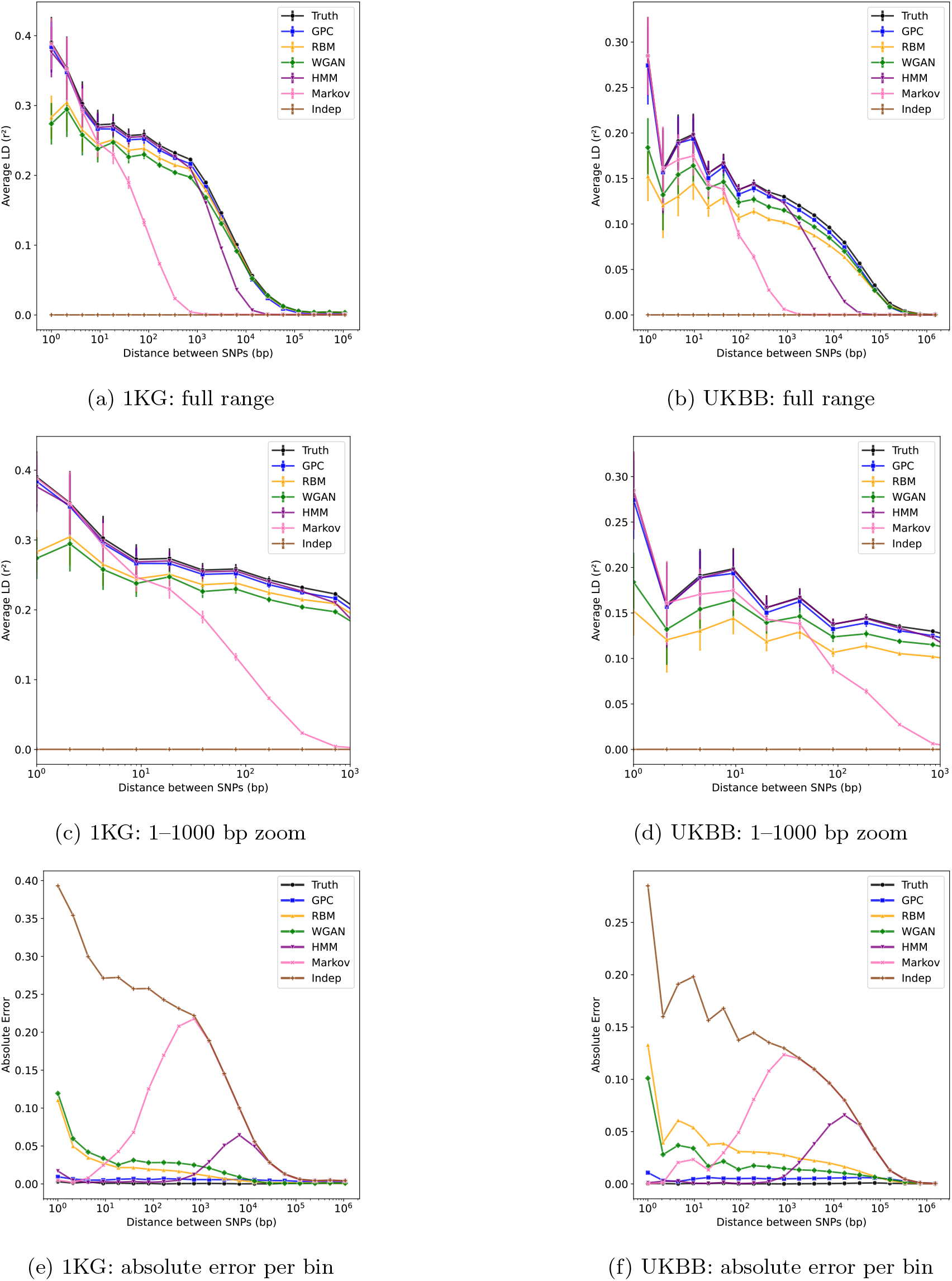
Patterns of linkage disequilibrium (LD) in the AGs generated by models. We compare LD betweens pairs of SNPs in the AGs generated by each of the models trained on the 1KG (left) and the UKBB (right) datasets respectively. We plot mean squared correlation *r*^2^ within bins of physical distance (error bars denote the standard error of the mean); truth represents the LD distribution of the training data. We plot the decay of LD across the full length scale of the locus (top row); at shorter length scales (1–1000 bp; middle row); and the absolute error in mean *r*^2^ per bin relative to the test data (bottom row) (see Table S2 for detailed metrics).

**Figure 4:**
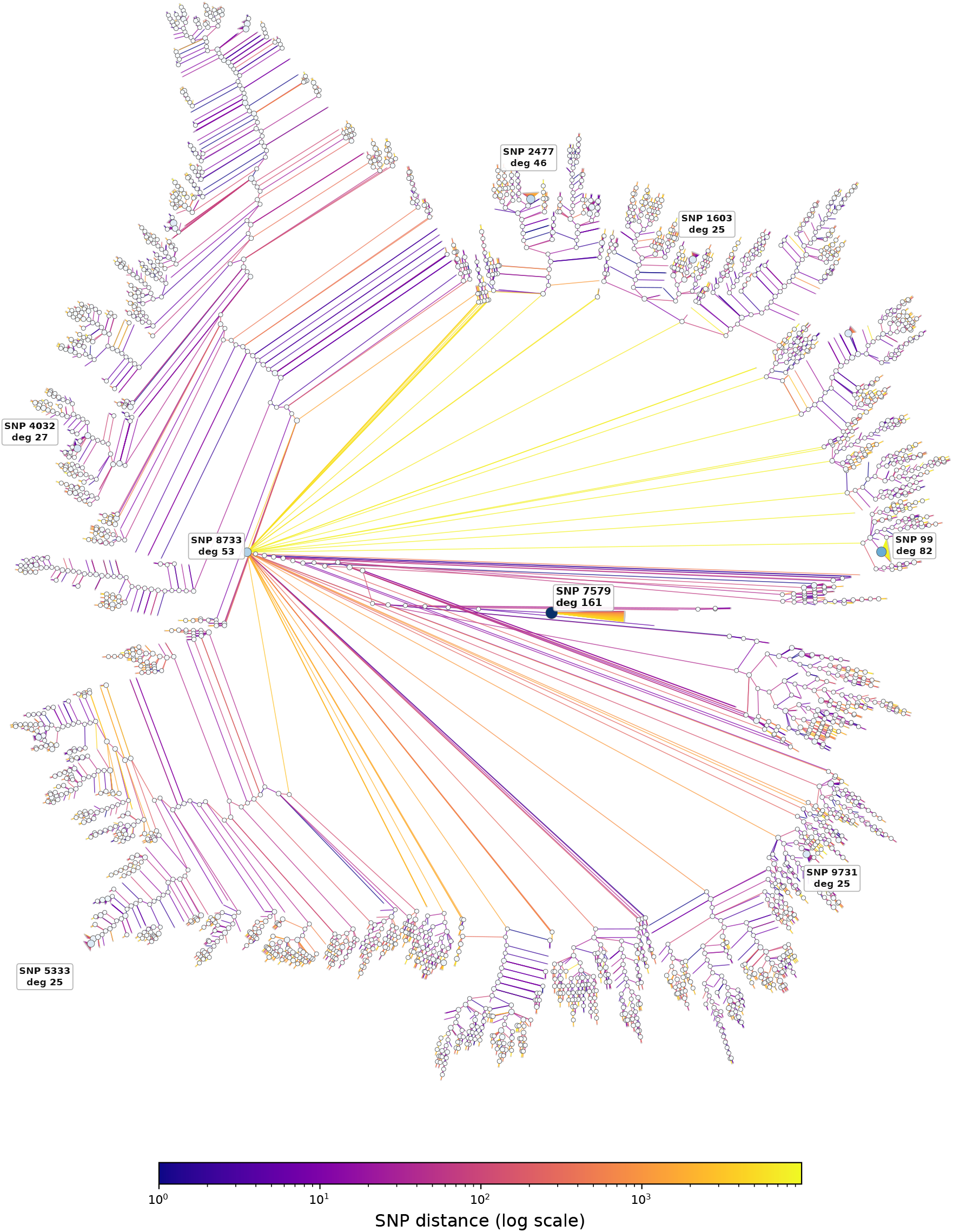
Compact radial visualization of the Chow–Liu Tree (CLT) learned from the 1KG dataset. Starting from the full 10,000-SNP tree (Supplementary Figure S4), all degree-2 nodes forming unbranched chains are contracted into single edges between their endpoints, retaining only branching points and leaves. Edge color encodes genomic distance |*i* − *j*| on a log scale (purple: nearby; yellow: distant). Node size scales with degree; the top 8 highest-degree nodes are labeled. Unlike an HMM, which forms a single unbranched chain, the CLT exhibits rich branching structure with long-range edges connecting SNPs thousands of positions apart, capturing distal LD directly rather than routing it through all intermediate variables.

The non-chain tree topology allows the model to place strongly correlated SNPs close together in the tree regardless of their genomic positions, enabling direct propagation of information across long ranges without passing through all intermediate variables.

### 2.6 GPC improves imputation accuracy across diverse scenarios

A key downstream application of generative models and AGs is genotype imputation. To assess the quality of artificial genomes (AGs) produced by GPC and competing generative models, we evaluate their utility in several imputation settings across the datasets described in Section 2.2. One approach is to generate AGs from these models, which then serve as reference panels for Impute5 [32] and other standard imputation tools. Alternatively, as described in Section 2.1, GPC can perform *direct imputation* by leveraging efficient conditional probability computation over the learned model, without the need to generate AGs.

Following the same training protocol as in Section 2.5, we use 80/20 train/test splits and generate the same number of AGs per dataset. For population-specific experiments, the same 80/20 split is applied within the target population. Following the experimental protocol established in prior work [32, 33, 44], we evaluate imputation accuracy at each SNP by removing the selected SNP from the test haplotypes while keeping all other SNPs observed. Imputation is then performed conditional on the remaining observed SNPs in the test haplotypes. We calculate the squared Pearson correlation (*r*^2^) between the imputed posterior probabilities of carrying allele 1 and the true allelic state at the target SNP (excluding monomorphic SNPs), aggregating results into bins by minor allele frequency (MAF) with 95% confidence intervals computed from 10 bootstrapped replicates. This imputation protocol follows the approach of Yu et al. [44] (Sections 2.6.1 and 2.6.2). We additionally evaluate imputation from genotyping arrays following Rubinacci et al. [32] and Browning et al. [33] for which we use the high-coverage 1KG dataset (Section 2.6.3) (additional details in Section 4.3).

We consider two broad scenarios for imputation. In the first (general) scenario, imputation is applied to genotypes from a cosmopolitan sample (test set) consisting of individuals from multiple ancestries with AGs generated by different models that were also trained on a cosmopolitan panel of matched ancestries consisting of a distinct set of individuals (training set). This setting reflects common practice in large-scale genetic studies where diverse reference panels are used to impute missing genotypes across heterogeneous cohorts. The second (population-specific) scenario considers a setting in which imputation is to be performed in a population for which reference genomes are limited or restricted. Because the target population might be under-represented in public reference panels, imputation accuracy in these populations might be adversely impacted. This scenario is particularly relevant for underrepresented populations in genomic research, where privacy constraints or limited sequencing resources may restrict the availability of ancestry-matched reference data. In a realistic setting, a researcher may only have access to publicly available European reference genomes while population-specific data remains private. GPC and other generative approaches can learn from such private data and either generate AGs or perform direct imputation without exposing the underlying genomes. For the population-specific analyses, we designate a target population which is limited or restricted in access. We consider two sets of target populations: all individuals of non-European ancestry and all individuals of African ancestry. In each of these settings, we also explore how the accuracy of imputation is impacted when AGs are combined with real genomes (possibly with different genetic ancestry characteristics than the target population) that might be available.

#### 2.6.1 General Imputation

We first examine the general setting, in which models are trained on genomes from diverse ancestries and are then used to impute into target genomes drawn from similar ancestries. To mimic this scenario, we train and test models on random splits of the 1KG and UKBB datasets. For 1KG, we also include a comparison to 5,008 AGs based on fitting WGAN and RBM to the same genomic region that were made available by the authors of [25].

We see that reference panels composed entirely of real genomes used with Impute5 achieve the highest imputation accuracy across MAF bins, providing an approximate upper bound on achievable performance (Figure 5a for 1KG and Figure 5b for UKBB). Among the generative models, GPC obtains the highest imputation accuracy across MAF bins. Averaged across both datasets, GPC (direct) achieves a 0.168 (27.5%) improvement in *r*^2^ over the next best method, RBM (0.230 (174%) for low-frequency variants with MAF *<* 1%). GPC (direct) imputation, via conditional probability computation rather than simulated AGs, tends to be the more accurate version, achieving a 0.105 (15.5%) improvement in *r*^2^ (0.195 (61.5%) for low-frequency variants) over GPC with AG-based imputation, likely due to directly targeting the imputation objective rather than relying on simulated AGs. To assess the robustness of these findings, we repeated the UKBB general imputation experiment on five additional LD blocks across distinct chromosomes (see Section 4.1.2 for details) to find the same qualitative pattern holds across all regions: GPC (direct) consistently improves over RBM (the next best method) across the range of MAF (Figure 5c; additional per-chromosome results in Supplementary Figure S5).

**Figure 5:**
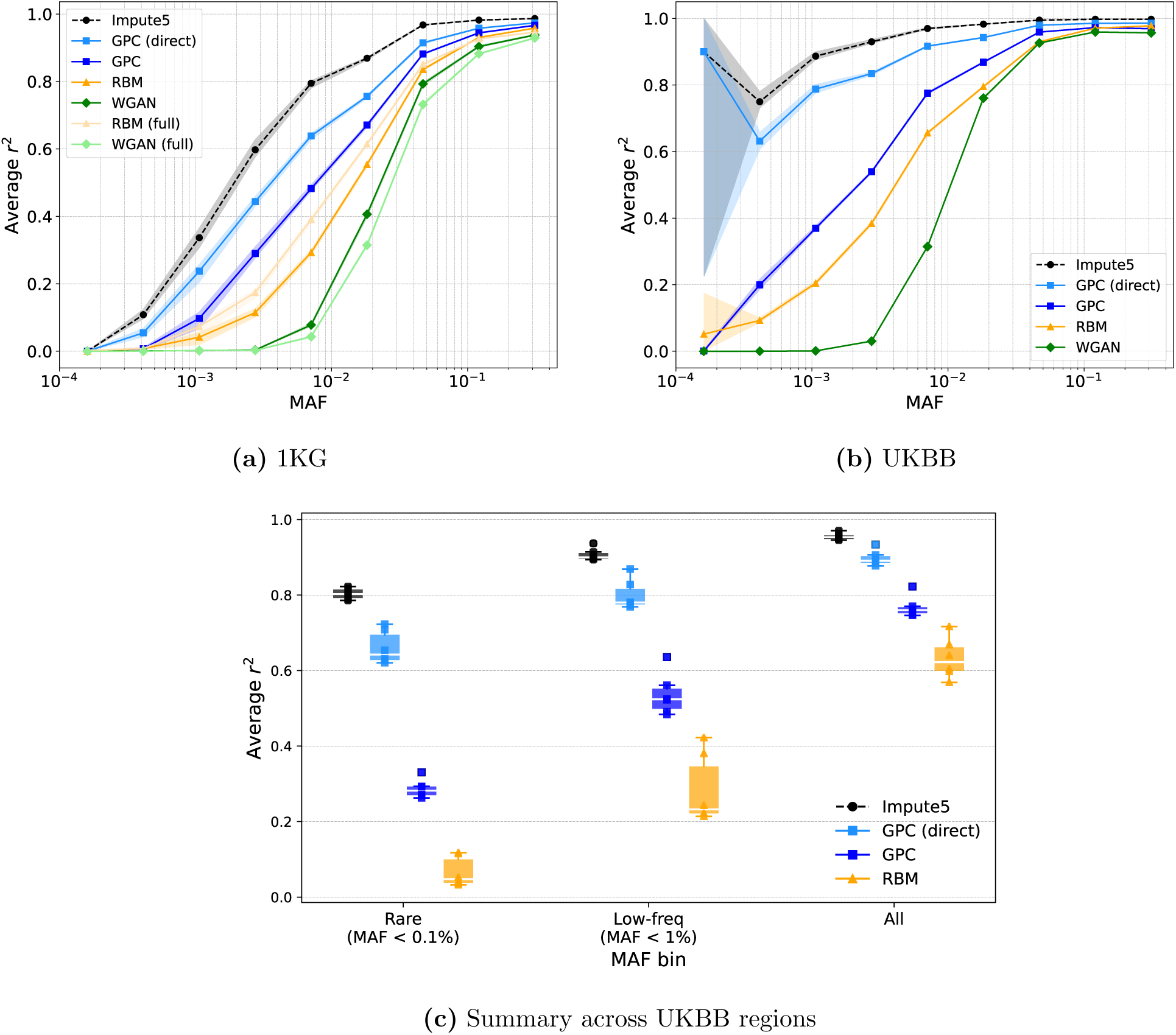
General imputation. The light blue lines denote direct imputation using GPC while the other lines show the results of imputation using Impute5 with AG reference panels in the 1KG (left) and the UKBB (right) datasets respectively. We also plot the results of imputation using real reference genomes (black dotted lines) while RBM/WGAN (full) show results using AGs provided by the authors of [25] in their analysis of the same locus in the 1KG dataset (see Table S3 for detailed metrics). (c) summarizes UKBB imputation performance for the best-performing methods (excludes WGAN) by MAF bin, averaged across chr22 (the main UKBB region shown in (b)) and five additional chromosomes (chr2, 11, 12, 14, 18; see Supplementary Figure S5), demonstrating that results are robust across genomic regions.

#### 2.6.2 Population-Specific Imputation

We next consider imputation in populations for which reference genomes are limited or restricted. Because large public datasets are predominantly of European ancestry, we consider scenarios where the target population consists of individuals of either non-European or African ancestry. In this setting, imputation accuracy in the target population can be degraded when public European reference genomes are the only data available to the imputation method. AGs specific to the target population generated by GPC can mitigate this challenge by learning from an ancestry-matched set of private reference genomes. We also evaluate combined reference panels formed by augmenting real European data with population-specific AGs.

Analogous to the general imputation experiments, GPC (direct) remains the most accurate across MAF bins (Figures 6 and S6 for 1KG and UKBB respectively). Averaged across datasets and ancestries, GPC (direct) achieves a 0.154 (33%) improvement in *r*^2^ over the next best method, RBM (0.202 (279%) for low-frequency variants). Unlike in the general imputation setting, GPC, when used for direct imputation, tends to be more accurate than using Impute5 with a reference panel of European genomes, likely due to the distributional mismatch between the reference and target populations. This holds consistently across the 1KG dataset. In the UKBB dataset, GPC (direct) outperforms Impute5 for common variants but is slightly less accurate for rare variants. We attribute this to our balanced sampling strategy (Section 4.1.2), in which the European reference panel available to Impute5 and the population-specific training data for the generative models are matched in sample size. Under this design, a large European reference panel (even one mismatched in ancestry) can partially compensate for distributional mismatch at rare variants, where population-specific sample sizes are most limiting. Importantly, when the real European genomes are incorporated into the training of the generative models (the “combined” setting), all methods improve, and GPC (direct combined) remains competitive with or above Impute5 across MAF bins, indicating that the gains from ancestry matching and from larger sample size are complementary. Averaged across datasets and ancestries, GPC (direct) achieves a 0.056 (12.3%) improvement in *r*^2^ over Impute5 using European genomes (0.012 (42.1%) for low-frequency variants). Finally, we find that combining population-specific AGs with European genomes consistently improves accuracy across all MAF bins. Averaged across datasets and ancestries, GPC (direct) achieves a 0.011 (1.8%) improvement in *r*^2^ over itself when including European genomes in the reference panel (0.023 (12.3%) for low-frequency variants).

**Figure 6:**
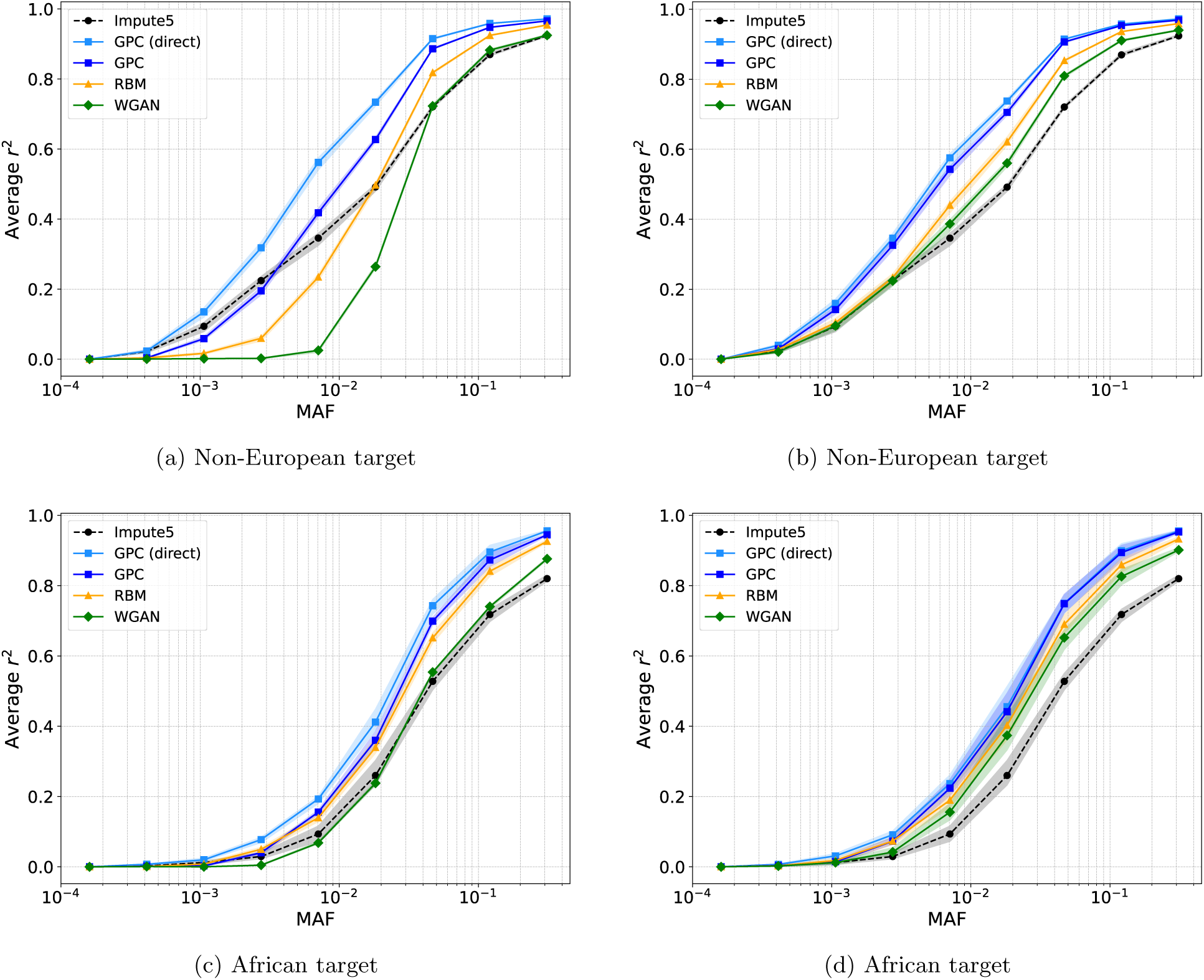
Population-specific imputation in the 1KG dataset. On the left, the AG lines show imputation results using population-specific AGs (where the target genomes correspond to non-European populations in part a and African populations in part c) while the light blue line shows direct imputation with GPC trained on population-specific genomes. On the right, the AG lines show results using combined reference panels (real European genomes plus population-specific AGs) while the light blue line shows direct imputation with GPC trained on both European and population-specific genomes (where the target genomes correspond to non-European populations in part b and African populations in part d). The black line shows results using Impute5 with real European genomes as the reference panel (see Table S5 for detailed metrics).

Taken together, these results demonstrate that population-specific AGs from GPC can substantially enhance imputation accuracy in underrepresented groups, either by augmenting existing reference panels or through direct conditional imputation. The magnitude of improvement depends on several interacting factors: the degree of genetic distance between the target population and the available public reference panel, the sample size of the private population-specific data, and the frequency of the variants being imputed. When public reference data is both ancestrally mismatched and matched in size to the private data, as in our balanced UKBB design, gains over a large European-only panel are most pronounced for common variants, while rare variant performance remains sensitive to total sample size regardless of ancestry. These findings suggest that in practice, combining ancestry-matched private data with available public genome (even those from a different ancestry) will generally yield the best imputation accuracy, and that the relative value of each component will depend on the specific target population and variant frequency spectrum.

#### 2.6.3 Array-based Imputation

We also evaluate GPC in a more realistic scenario of imputation from variants genotyped on a commonly-used SNP array. In this experiment, we attempted to impute SNPs genotyped in the high-coverage 1KG dataset from variants that were typed on the HumanOmni5Exome array (see Section 4.1.3 for details).

GPC (direct) remains the most accurate in both the general (Figure 7) and population-specific settings (Figure 8). In the general setting, GPC (direct) achieves a 0.020 (3.5%) improvement in *r*^2^ over the next best method, RBM (0.043 (14.3%) for low-frequency variants). In the population-specific setting, averaged across both ancestries, GPC (direct) achieves a 0.030 (5.3%) improvement in *r*^2^ over RBM (0.040 (12.7%) for low-frequency variants). Similar to the previous population-specific setting, GPC (direct) outperforms Impute5 with the European-only reference panel, achieving a 0.285 (96.5%) improvement in *r*^2^ averaged across both ancestries (0.324 (1,123%) for low-frequency variants). In contrast to our previous imputation experiments, here the best performance is achieved by GPC (direct) trained solely on the population-specific data, rather than a combined population-specific plus European dataset. Thus, the optimal imputation strategy is likely determined by a combination of genetic ancestries, sample size, and SNP sets, and needs further investigation. These results suggest that in realistic imputation scenarios, population-specific modeling may provide accuracy gains over approaches that aggregate across populations.

**Figure 7:**
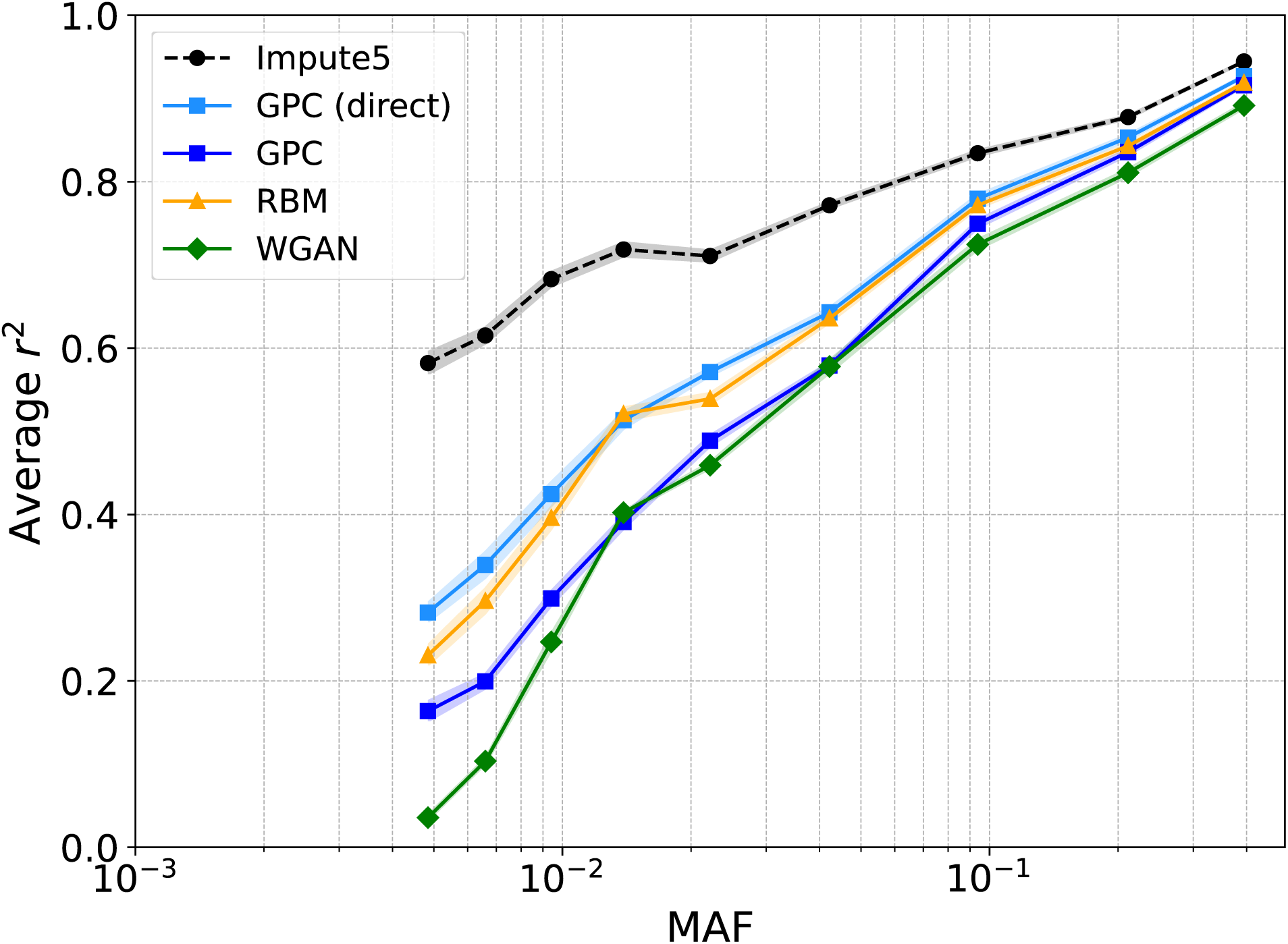
Array-based imputation. This task aims to impute SNPs from the HumanOmni5Exome array using high-coverage 1KG data. The black and light blue lines show results using Impute5 with real training data as the reference panel and direct imputation using GPC, respectively. The other lines show results using Impute5 with AG reference panels (see Table S7 for detailed metrics).

**Figure 8:**
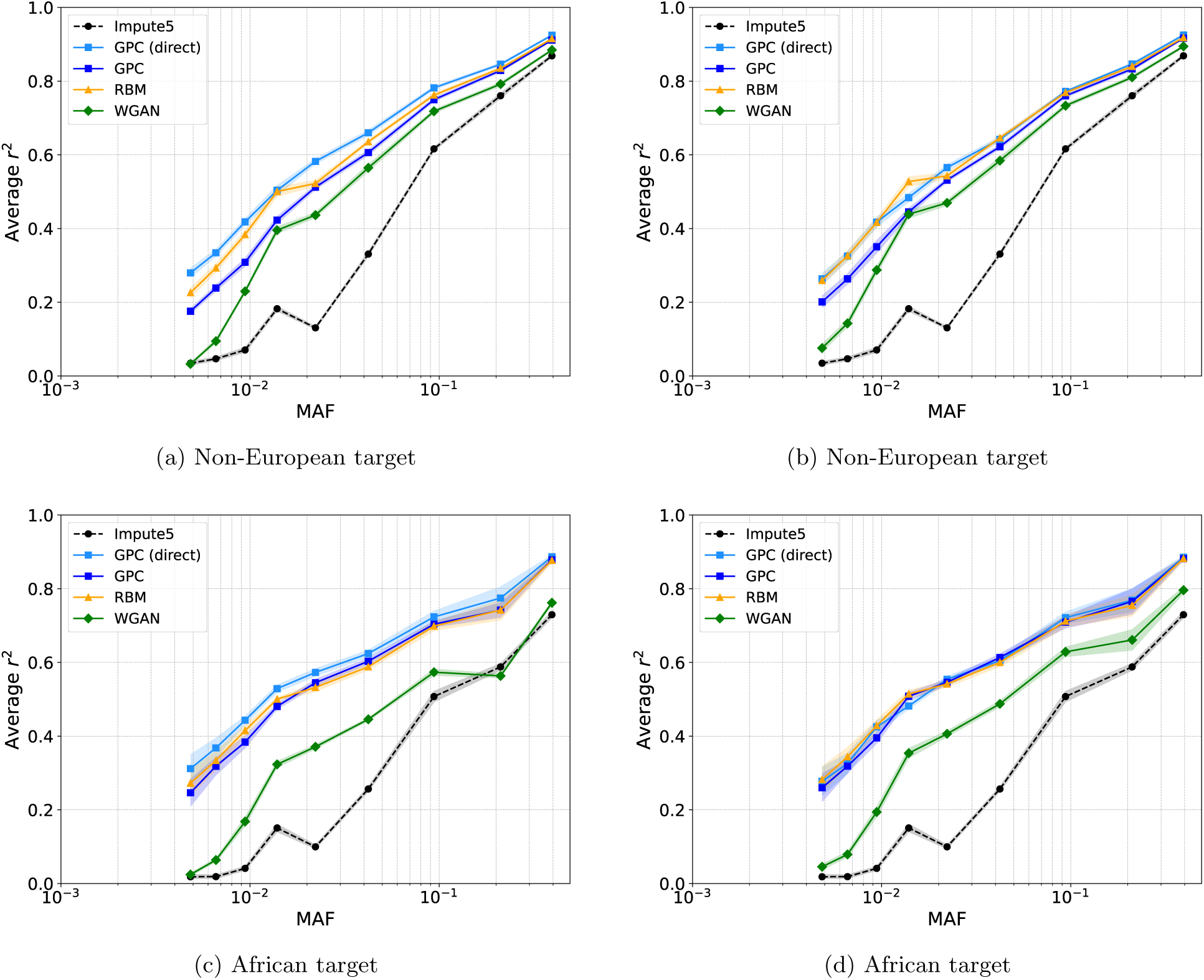
Array-based imputation (population-specific). The black line shows results using Impute5 with real European reference genomes. On the left (parts a and c), the AG lines show results using population-specific AGs alone, and the light blue line shows direct imputation with GPC trained on population-specific data. On the right (parts b and d), the AG lines show results using combined reference panels (real European data plus population-specific AGs), and the light blue line shows direct imputation with GPC trained on both European and population-specific data (see Table S8 for detailed metrics).

### 2.7 GPC balances considerations of privacy and utility

To evaluate the privacy of the AGs, we compute the nearest neighbor adversarial accuracy (*AATS*) metric introduced by Yale et al. [45]. The metric decomposes into two components: *AA*_TRUTH_, which measures whether real samples are closer to other real samples than to synthetic ones, and *AA*_SYN_, which measures the analogous property for synthetic samples. Values near 0.5 for both components indicate an ideal balance between utility and privacy. We report *AA*_TRUTH_ and *AA*_SYN_ separately rather than their average, as the overall *AATS* can mask pathological cases where poor values in one component cancel those in the other. We evaluate on both training and held-out test sets and standardize genotypes prior to distance computation as this upweights the impact of rare variants which tend to be more challenging to model (see Section 4.5.1 for details).

Under this evaluation, GPC achieves *AA*_SYN_ closest to 0.5 across all splits and both datasets, and *AA*_TRUTH_ closest to 0.5 in all but one case (1KG test, where WGAN is marginally closer: 0.818 vs. 0.820), indicating the best overall balance between utility and privacy (Table 2). The near-zero *AA*_SYN_ values for RBM paired with near-unity *AA*_TRUTH_ reveal a distinct failure mode: real samples still find other real samples as their nearest neighbors (real-to-real distances are smaller than real-to-synthetic), but each synthetic sample is closer to some individual real genome than to any other synthetic sample. Rather than forming a coherent distribution, RBM produces samples that float near individual training haplotypes without capturing broader structure, a privacy risk because each synthetic genome can potentially be traced back to a specific training individual. Conversely, WGAN exhibits uniformly high values for both components, indicating that its synthetic distribution occupies a region largely disjoint from the real data, sacrificing utility. We note that even for GPC, *AA*_TRUTH_ remains noticeably above 0.5, suggesting that the AGs are still somewhat distinguishable from real data.

**Table 2:** Nearest neighbor adversarial accuracy (*AA*_TRUTH_, *AA*_SYN_) for AGs generated by RBM, WGAN, and GPC, evaluated on training and test splits from 1KG and UKBB. Distances are standardized Euclidean. This upweights rare variants relative to common ones, exposing failures in rare-variant modeling. Values closest to 0.5 are bolded.

| Dataset | Metric | RBM | WGAN | GPC |
| --- | --- | --- | --- | --- |
| **1KG** | $AA_{\text{TRUTH}}$ (Train) | 0.9561 | 0.8103 | **0.7185** |
| | $AA_{\text{TRUTH}}$ (Test) | 0.9916 | **0.8183** | 0.8199 |
| | $AA_{\text{SYN}}$ (Train) | 0.0024 | 0.7356 | **0.4225** |
| | $AA_{\text{SYN}}$ (Test) | 0.0212 | 0.7680 | **0.4912** |
| **UKBB** | $AA_{\text{TRUTH}}$ (Train) | 0.9934 | 0.9664 | **0.9096** |
| | $AA_{\text{TRUTH}}$ (Test) | 0.9970 | 0.9692 | **0.9240** |
| | $AA_{\text{SYN}}$ (Train) | 0.0094 | 0.7828 | **0.5380** |
| | $AA_{\text{SYN}}$ (Test) | 0.0114 | 0.7730 | **0.4686** |

## 3 Discussion

In this work, we introduced GPC, a deep generative model for genetic variation based on hidden Chow-Liu trees (HCLTs) and probabilistic circuits (PCs). Unlike prior deep generative models (e.g., GANs, VAEs, and RBMs), GPC combines tractable probabilistic inference with an expressive latent variable structure, enabling principled convergence monitoring via held-out log-likelihood, accurate reconstruction of population structure and linkage disequilibrium patterns across all distance scales, and direct genotype imputation through exact conditional probability computation. In population-specific settings, GPC (direct) consistently outperforms other deep generative methods as well as Impute5 using publicly available European reference genomes, with particularly strong gains for low-frequency variants that are disproportionately population-specific. We also found that GPC AGs provide improved privacy properties compared to RBMs and WGANs, though we emphasize that demonstrating improved performance on a single metric does not guarantee complete privacy preservation. Other attack models, such as membership inference attacks [46, 47] or attribute inference attacks [48], may still pose risks. Future work should explore whether GPC can be extended to provide formal privacy guarantees through differentially private EM algorithms [49].

Despite these advantages, several limitations remain. First, scaling GPC to full genome length remains challenging and will likely require hierarchical or distributed approaches. A straightforward approach to whole genome data would involve applying GPC to LD blocks which are treated as independentof each other though such an approach is likely to ignore LD across blocks. Second, our current implementation of GPC is restricted to haploid genomes to enable direct comparisons with prior approaches. Extending GPC to generate diploid genotype data would increase the applicability of the model. Third, like all generative approaches, GPC inherits biases from the training data, and ensuring fairness across diverse cohorts remains an open challenge. Fourth, additional downstream benchmarks such as rare variant association studies, fine-mapping, or polygenic risk scores would provide a more complete picture of utility. Finally, our experiments focused on single genomic regions; genome-wide evaluation would better assess robustness across varying LD structures and recombination rates.

Additionally, recent work has extended the space of deep generative approaches for genetic data to include VAEs [29, 30], GANs operating in reduced-dimensional spaces [26, 28], and diffusion models [31], as well as genome-wide WGAN and CRBM variants [25]. Systematically comparing GPC against these approaches across imputation and other downstream tasks, and extending all methods to genome-wide scales, remains an important direction for future work.

## 4 Materials and Methods

### 4.1 Datasets

#### 4.1.1 1000 Genomes Project Phase 3 (1KG)

We first use data from the 1000 Genomes Project (Phase 3), consisting of 2,504 unrelated diploid genomes spanning diverse ancestries. Because the genomes are phased, we split each into its two constituent haplotypes, yielding 5,008 haplotypes in total. To avoid any possibility of data leakage, all train/test splits are performed at the diploid-individual level, before haplotype separation.

Following prior work [24, 25], we analyze a contiguous 10K-SNP region on chromosome 15 (15:27,379,578 – 15:29,625,035). This dataset is used primarily to compare GPC to baselines and to evaluate how well it captures population and LD structure. It is also used in our single-SNP imputation experiments.

#### 4.1.2 UK Biobank (UKBB)

To evaluate performance in a larger, secondary cohort, we analyze genotypes from the UK Biobank (UKBB). We analyzed high-quality imputed SNPs (with a hard call threshold of 0.2 and an INFO score ≥ 0.8) with MAF ≥ 0.1%. We apply standard quality control by retaining SNPs that are under Hardy-Weinberg equilibrium (*p<* 10^−6^) and are confidently imputed in more than 99% of the individuals. We retain individuals with no kinship to other individuals (UKBB field 22021) and exclude those with missing ancestry information. Following these procedures, we select an LD block on chromosome 22 (22:29,456,546 – 22:32,665,772). This yields 337,862 individuals and 9,820 SNPs. To assess robustness across genomic regions, we additionally select five LD blocks on separate chromosomes: chromosome 2 (2:118,367,466 – 2:121,303,783; 9,227 SNPs), chromosome 11 (11:101,331,121 – 11:103,959,636; 9,507 SNPs), chromosome 12 (12:85,990,426 – 12:89,682,122; 9,823 SNPs), chromosome 14 (14:72,889,615 – 14:76,444,767; 9,248 SNPs), and chromosome 18 (18:51,554,175 – 18:55,213,838; 9,415 SNPs). These additional regions are used for the general imputation replication experiment. We phased all data using Beagle 5.5.

To ensure computational feasibility for all methods while maintaining statistical power, we randomly select 5,000 individuals each from the European (EUR), non-European (Non-EUR), and African (AFR) ancestry groups, yielding 10,000 haplotypes per group after phasing. Due to partial overlap between the Non-EUR and AFR subsets, the combined imputation dataset includes 26,924 total haplotypes. Similar to 1KG, we use this dataset to evaluate population/LD structure and single-SNP imputation. The balanced sampling strategy also allows us to assess whether equal amounts of European and non-European data can compensate for distributional mismatch when imputing into non-European test populations.

#### 4.1.3 High-coverage 1KG

To evaluate GPC in a realistic imputation scenario, we consider the high-coverage whole-genome sequencing release of 1KG, using the same genomic region as in the earlier 1KG analysis, mapped to build 38 coordinates (15:27,134,431 – 15:29,332,831). We restrict the analysis to the same set of 2,504 unrelated individuals. This region contains 56,766 variants. After selecting biallelic SNPs and removing those with minor allele count (MAC) ≤ 20, 14,670 SNPs remain.

Among these, 2,119 SNPs are present on the HumanOmni5Exome-4v1-2 A genotyping array. The remaining 12,551 SNPs (86%) are imputed simultaneously, providing a setting that closely mirrors real imputation pipelines. This dataset is used for our array-based imputation experiments, both overall and stratified by ancestry.

### 4.2 Model and Inference Methods

#### 4.2.1 Hidden Chow-Liu Trees

Hidden Chow-Liu trees (HCLTs) [35] represent a distribution over a collection of random variables (RVs) (**Z** = (*Z*_1_,…, *Z*_*N*_), **X** = (*X*_1_,…, *X*_*N*_)). **Z** denotes hidden or latent RVs while **X** denotes observed RVs. The joint distribution is described by a graphical model (𝒢) in which the nodes in the graph represent the

RVs and the lack of edges among the nodes represents conditional independence assumptions. In HCLTs, we have edges from each hidden variable to its corresponding observed random variable (*Z*_*n*_ → *X*_*n*_) while the edges among the hidden variables form a tree. When the graph over the hidden variables is a chain (*Z*_1_ → *Z*_2_ → … *Z*_*N*_), we obtain a hidden Markov model (HMM). By permitting tree-structured graphs, HCLTs generalize HMMs and can provide a better representation of the data to capture long-range dependence, *e*.*g*., RV *X*_2_ and *X*_6_ are highly correlated without being correlated with *X*_3_, *X*_4_, *X*_5_ (Figure 1).

For genetic variation data over *N* single nucleotide polymorphisms (SNPs), each *X*_*n*_ denotes the genotype value at SNP *n* ∈ {1,…,*N* } (*X*_*n*_ ∈ {0, 1} when we model haploid genomes). Each *Z*_*n*_ is a discrete RV that can take one of *L* values (*Z*_*n*_ ∈ {0,…,*L* − 1}). Figure 1 demonstrates how to construct an HCLT using genetic data. Given a dataset 𝒟 that contains 6 SNPs (Figure 1(a)), we first invoke the Chow-Liu algorithm to generate a tree over the latent variables associated with each SNP (Figure 1(b)). The Chow-Liu algorithm computes pairwise mutual information between all variable pairs and finds the maximum-weight spanning tree, which can be done efficiently even for large SNP panels since it only requires computing pairwise statistics and solving a minimum spanning tree problem. The tree encodes strong variable dependencies by placing highly correlated SNPs (e.g., *X*_1_ and *X*_3_) closer in the generated tree. Finally, the HCLT is constructed by adding an edge from every latent variable *Z*_*i*_ to its corresponding observed variable *X*_*i*_ (Figure 1(c)). The parameters of the HCLT are those associated with the discrete conditional probability distributions *P* (*X*_*n*_|*Z*_*n*_) and *P* (*Z*_*n*_|*Z*_Pa(*n*)_), where Pa(*n*) denotes the parent of node *n* in the tree.

#### 4.2.2 Probabilistic Circuits

Probabilistic Circuits (PCs) [37, 38] are a class of probabilistic models that support tractable probabilistic inference. These capabilities have allowed PCs to perform various probabilistic reasoning tasks that are out of reach for most deep generative models [50, 51], including problems in explainable AI [52, 53, 54], algorithmic fairness [55, 56], and missing data robustness [57, 54, 58]. PCs are furthermore appealing for their expressive power and suitability for density estimation, with recent advances in structure learning [59] and parameter estimation [56, 35] allowing PCs to accurately capture useful correlations in the data.

##### Representation

PCs are an umbrella term for a wide family of tractable probabilistic models [60], including arithmetic circuits [61], sum-product networks [62], and cutset networks [63]. A PC (𝒢, ***θ***) represents a joint probability distribution Pr(**X**) over random variables **X** through a directed acyclic graph (DAG) 𝒢 parametrized by ***θ***. The DAG 𝒢 consists of three types of nodes: *input, sum*, and *product*. Each leaf node is an input node; each inner node *n* (i.e., sum or product) receives inputs from its children `ch`(*n*). Each node *n* ∈ 𝒢 encodes a probability distribution Pr_*n*_, which is defined recursively as follows:

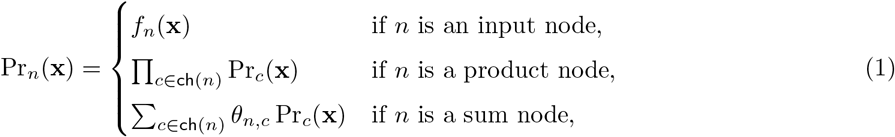

where *f*_*n*_(**x**) is a univariate input distribution (e.g., Binomial, Gaussian), and *θ*_*n,c*_ denotes the parameter that corresponds to edge (*n, c*). Intuitively, a product node defines a factorized distribution over its inputs, and a sum node represents a mixture over its input distributions weighted by *θ*. Finally, the probability distribution of a PC is defined as the distribution represented by its root node. The size of a PC (𝒢, ***θ***) is defined as the number of parameterized edges in its DAG 𝒢.

##### Inference

In contrast to many other generative models, PCs support efficient reasoning over their encoded distribution. One can compute likelihoods by evaluating the PC feed-forward as in Equation 1. Many common reasoning tasks such as marginal probabilities and maximum a posteriori probability (MAP) are also supported by PCs. To guarantee the efficiency for computing these queries, the DAG of the PC should satisfy certain structural constraints. Please refer to [64] for a more detailed summary of various inference scenarios for PCs.

To support linear-time computation (with respect to the size of the PC) of arbitrary marginal queries, PCs need to satisfy two structural properties: smoothness and decomposability. Both are properties of the scope *ϕ*(*n*) of PC units *n*, that is, the collection of variables defined by all its input nodes.

###### Definition 1

**(Smoothness)** *A PC* (𝒢, ***θ***) *is smooth if for any sum node n* ∈ 𝒢, *its children have identical scope:* ∀*c*_1_, *c*_2_ ∈ `ch`(*n*) : *ϕ*(*c*_1_) = *ϕ*(*c*_2_).

###### Definition 2

**(Decomposability)** *A PC* (𝒢, ***θ***) *is decomposable if for any product node n* ∈ 𝒢, *its children have disjoint scopes:* ∀*c*_1_, *c*_2_ ∈ `ch`(*n*), *c*_1_ ≠ *c*_2_ : *ϕ*(*c*_1_) ∩ *ϕ*(*c*_2_) = ∅.

Given a smooth and decomposable PC, querying an arbitrary marginal probability boils down to a feedforward evaluation of its DAG, thus the computation time is linear with respect to the size of the PC.

#### Sampling and Conditional Inference

Sampling from a PC is performed via ancestral sampling. Starting from the root node, we traverse sum nodes by sampling a child proportionally to the edge weights *θ*_*n,c*_, and traverse product nodes by recursively sampling from all children (since they define independent factorizations over disjoint variable sets). At input nodes, we sample from the corresponding univariate distributions. This procedure generates a complete assignment to all variables in time linear in the circuit size.

For genotype imputation, we compute conditional probabilities *p*(*X*_missing_|*X*_observed_). Smooth and decomposable PCs support exact marginalization over any subset of variables: to marginalize out a variable *X*_*i*_, we simply replace its input distribution with the constant 1 (summing over all possible values). Conditional queries are then computed as ratios of two marginal queries [37]:

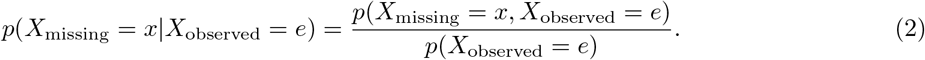

Both the numerator and denominator can be computed via a single feedforward pass through the circuit, making conditional inference efficient.

#### 4.2.3 HCLTs as Probabilistic Circuits

HCLTs can be represented as smooth and decomposable PCs, meaning they support efficient (i.e., linear in the size of the PC) computation of marginal queries and likelihoods. When considering both the observed and hidden variables, HCLTs are also deterministic (i.e., for any sum node, its children have disjoint support), meaning that the MAP instance arg max_**x**∈**X**,**z**∈**Z**_ Pr_*n*_(**x, z**) can be evaluated in linear time. However, HCLTs are not deterministic with respect to the observed variables **X** alone, which has implications for parameter estimation as described below.

#### 4.2.4 Parameter Estimation

Any probabilistic graphical model (PGM) can be transformed into a PC that encodes the same probability distribution. We demonstrate the high-level idea of this transformation, and refer interested readers to [38] for more details. To transform an HCLT into an equivalent PC, we iteratively encode every conditional probability Pr(**X**_*n*_|**X**_Pa(*n*)_) by representing each possible value of **X**_*n*_ as a sum node. Thus, the probability of Pr(**X**_*n*_ = **x**_*n*_|**X**_Pa(*n*)_ = **x**_Pa(*n*)_) can be represented by the weight of an edge connecting **x**_*n*_ and **x**_Pa(*n*)_. Take the HCLT in Figure 1 as an example. The conditional probabilities are encoded into a single PC in a bottom-up manner: we first encode Pr(*X*_5_|*Z*_5_) and then followed by Pr(*X*_4_|*Z*_4_) and Pr(*Z*_5_|*Z*_4_), and so on.

If a PC is smooth, decomposable, and deterministic, its maximum-likelihood estimation (MLE) parameters can be efficiently learned in closed form [65]. To formalize the MLE parameters, we define the context *γ*_*n*_ of any node *n* as follows. The context of the root node *n*_*r*_ is its support `supp`(*n*_*r*_). The context of any other node is the intersection of its support and the union of its parents’ contexts:

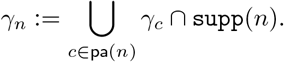

For any sum node *n* and its child *c*, the associated MLE parameter 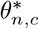 on a dataset 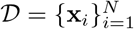 is

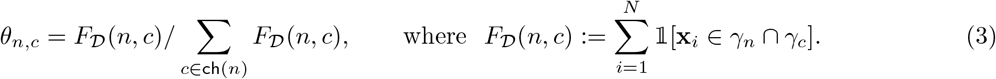

The quantity *F*_𝒟_(*n, c*) is called the *circuit flow* of edge (*n, c*). Intuitively, circuit flows count the number of samples in 𝒟 that “activate” an edge.

However, since HCLTs are not deterministic with respect to the observed variables **X**, MLE does not have a closed-form expression, and we instead resort to Expectation-Maximization (EM), where in the E step we compute the *expected circuit flow* given incomplete data, and in the M step we estimate the closed-form MLE parameters given those expected flows [66, 56].

Concretely, given a deterministic PC (𝒢, ***θ***) with root node *r* and an incomplete dataset 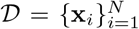, the parameters for the next EM iteration are given by [56]:

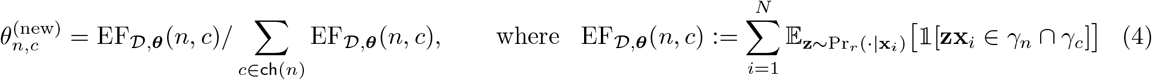

defines an expected version of circuit flow for edge (*n, c*) given samples with missing values in 𝒟.

Using their equivalent PC representations, HCLTs can be trained efficiently using the PC package PyJuice [40]. By representing graphical models (e.g., HCLTs) as PCs, we take advantage of the structure of the model to extensively parallelize the computation required by EM updates (Equation 4). Further, we develop specialized GPU kernels to significantly speed up the EM algorithm. As a result, despite the model having over 88 million parameters, a single EM epoch on the 1KG dataset can be done in less than 2 seconds.

### 4.3 Impute5 Specifications

Haploid imputation is specified in Impute5, with all other SNPs in the region serving as buffer (observed context). Further, we provide the corresponding fine-scale human recombination map for the chromosome being imputed. These are linked in the Impute5 documentation: https://github.com/odelaneau/shapeit4/tree/master/maps.

### 4.4 Training Specifications

Training GPC is straightforward, requiring minimal hyperparameter tuning. We set the number of latent states (for each hidden variable in the HCLT) to 128 for both datasets – the maximum feasible given memory constraints. We observe that changing the pseudocount parameter (used to smooth probability estimates) does not significantly impact the training procedure and can be left at a small default value (0.005).

Based on initial experiments with small validation sets, we find that training for 2,000–5,000 epochs works well for these datasets, though longer training is possible without overfitting (with minimal performance gains). For the 1KG data, we trained GPC for 5,000 epochs. Due to the larger sample size (and longer training time per epoch) in the UKBB data, we stopped at 2,000 epochs. Although we do not tune the pseudocount, this would be the only parameter to consider adjusting for further optimization. Additionally, since we use full-batch EM to learn the circuit parameters, we do not need to tune other standard hyperparameters such as learning rate or batch size.

A key advantage of GPC over other deep generative approaches is that we can probabilistically determine convergence by monitoring held-out log-likelihood, which provides an objective, quantitative stopping criterion. This stands in contrast to methods that lack tractable likelihoods, where convergence must be assessed through slower, less consistent visual inspection methods that are more prone to human error.

The baseline RBM and WGAN methods require substantially more time and experimentation to tune, primarily due to their larger hyperparameter spaces and inability to calculate likelihoods or determine convergence probabilistically. For RBMs, the partition function is intractable due to an exponentially large number of configurations over the visible and hidden nodes. Therefore, we must rely on indirect metrics such as Nearest Neighbor Adversarial Accuracy (AATS) [45] and visual overlap in principal component space. Similarly, WGANs do not define a probability distribution over the data, preventing direct evaluation of sample likelihoods. Here too, we must rely on visual overlap in the principal component space to assess convergence, a subjective and time-consuming process.

We utilize the training code and notebooks provided by the authors on GitHub to tune both methods. Although we initially struggled to achieve optimal performance on the 1KG African data subset, we resolved this issue after consulting with the authors. For the RBM, we modified several parameters (e.g., higher learning rate, fewer epochs, fewer Gibbs sampling steps). For the WGAN, we increased the batch size. Otherwise, all other parameters were kept at their default values. When training on the UKBB data, we retained the default parameters; similar to the GPC case, we used fewer epochs due to longer training time but still achieved good overlap in the principal component space based on visual inspection.

The simpler PGM baselines, Indep and Markov, model the observed SNPs directly without latent variables. Indep treats each SNP as an independent Bernoulli random variable, estimating *P* (*X*_*n*_ = 1) separately for each site via its sample mean (calculated by a single pass over the training data). Markov is a first-order Markov chain: it models each SNP conditioned only on the immediately preceding one in genomic order, *P* (*X*_*n*_ | *X*_*n*−1_), and is likewise fit in closed form via maximum likelihood. Both models therefore require no iterative training. For HMM, we use the EM algorithm with random initialization. Similar to GPC, we set the number of latent states for HMM at 128 and train for 2,000–5,000 epochs depending on the dataset. Using these settings, the held-out log-likelihood converged effectively. We tried higher values of latent states for HMM but these all resulted in worse log-likelihoods and slower training overall.

### 4.5 Evaluation Metrics

#### 4.5.1 Nearest Neighbor Adversarial Accuracy

To evaluate the utility and privacy of the AGs, we compute the AATS metric introduced by Yale et al. [45]:

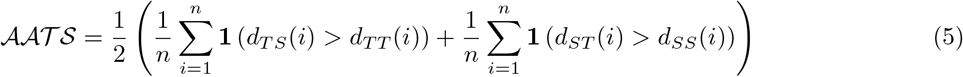

*T* represents the true (real) data and *S* represents the synthetic data. *d*_*TS*_(*i*) is the distance between sample *i* in the true data and its nearest neighbor in the synthetic data, while *d*_*TT*_ (*i*) is the distance between sample *i* in the true data and its nearest neighbor in the true data (not including itself). *d*_*ST*_ (*i*) and *d*_*SS*_(*i*) are defined analogously for the synthetic data. Following Yale et al. [45], we use Euclidean distance. The 𝒜𝒜𝒯𝒮 is an average of two components, which we denote *AA*_TRUTH_ and *AA*_SYN_:

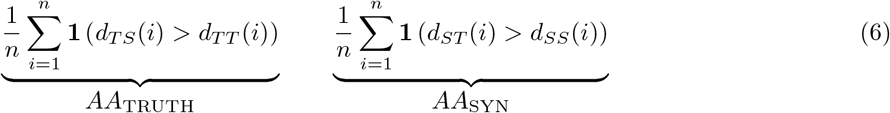

Each component returns a value between 0 and 1, with values near 0.5 indicating the ideal balance between utility and privacy. We do not report the overall 𝒜𝒜𝒯𝒮 because poor values in each component can cancel each other out: for instance, an *AA*_TRUTH_ of 0 paired with an *AA*_SYN_ of 1 would average to 0.5, masking both extreme memorization and a collapsed synthetic distribution.

##### Experimental setup

Using a 50/50 train/test split, we construct four datasets of equal size *n*: a real training set *R*_tr_, a real test set *R*_te_, and two disjoint halves of the generated synthetic data, *A*_1_ and *A*_2_. We compute the AATS components twice: pairing (*R*_tr_, *A*_1_) for train metrics and (*R*_te_, *A*_2_) for test metrics. For 1KG, *n* = 2,504, matching each real split exactly. For UKBB, the real dataset has 13,462 samples per split but we generated only 10,000 synthetic samples total, so we set *n* = 5,000: *A*_1_ and *A*_2_ are the first and second halves of the 10,000 generated samples, and 5,000 haplotypes are drawn without replacement from each real split using a fixed random seed.

##### Standardization

For each SNP, we compute the mean *μ*_*j*_ and standard deviation *σ*_*j*_ from *R*_tr_ only, and apply the transformation *x* ⟼ (*x* − *μ*_*j*_)*/σ*_*j*_ to all four datasets (*R*_tr_, *R*_te_, *A*_1_, *A*_2_) before computing any distances. Monomorphic sites (*σ*_*j*_ = 0) are left unchanged. Fitting the scaler on *R*_tr_ and applying it uniformly ensures that no information from the test set or the synthetic data influences the scaling. This choice is motivated by the fact that standardization reweights each SNP’s contribution to Euclidean distance by 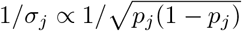, amplifying the contribution of rare variants (low MAF *p*_*j*_) and making the metric sensitive to failures in rare-variant modeling that would otherwise go undetected.

#### 4.5.2 Wasserstein Distance Calculation

To quantify differences between real and generated data, we used Wasserstein distances in two complementary settings. For comparisons in PCA space, we reported a two-dimensional Wasserstein distance computed between the empirical distributions of real (test set) and generated individuals. For each generative method, PCA was fit jointly on the real test samples and the corresponding generated samples (“coupled PCA”), and Wasserstein distances were evaluated in selected two-dimensional PCA subspaces (PC1–PC2, PC3–PC4, and PC5–PC6). Distances were computed using the entropically regularized optimal transport (Sinkhorn) algorithm as implemented in the ot Python package, with a regularization parameter set to *ε* = 2 *×* 10^−3^. Lower values indicate closer agreement between the distributions of real and generated samples in PCA space.

To assess similarity at the haplotype level, we computed one-dimensional Wasserstein distances between distributions of pairwise haplotypic distances. For each pair of haplotypes, we computed the Manhattan (cityblock) distance, which for haploid genomes equals the number of differing SNP positions, *i*.*e*., Hamming distance. We define two comparison metrics, both using the distribution of pairwise distances within the real test set (real–real) as the reference. The *within* metric compares internal diversity: it measures the Wasserstein distance between the distribution of all pairwise distances among AGs (generated–generated; *N*_*G*_(*N*_*G*_ − 1)*/*2 values for *N*_*G*_ AGs) and the distribution among real test haplotypes (real–real; *N*_*R*_(*N*_*R*_ − *1) /*2 values for *N*_*R*_ real genomes). This quantifies whether generated samples exhibit similar diversity to real samples. The *between* metric compares the distribution of cross-distances between generated and real test haplotypes (generated–real; *N*_*G*_ *× N*_*R*_ values) to the same real–real reference distribution. This quantifies whether generated haplotypes are as close to real haplotypes as real haplotypes are to each other. One-dimensional Wasserstein distances were computed using `scipy.stats.wasserstein_distance`, which handles distributions with different numbers of samples by treating them as empirical distributions.

## Acknowledgements

This work was supported, in part, by NIH grant GM153406, by National University of Singapore under its Start-up Grant SUG-251RES250, by the DARPA ANSR, CODORD, and SAFRON programs under awards FA8750-23-2-0004, HR00112590089, and HR00112530141, NSF grant IIS1943641, and gifts from Adobe Research, Cisco Research, Qualcomm, and Amazon. Approved for public release; distribution is unlimited. This research was conducted using the UK Biobank Resource under application 331277. We thank the participants of UK Biobank for making this work possible. We also thank the authors of [25], Burak Yelmen and Aurélien Decelle, for their assistance with training and parameter selection for the WGAN and RBM models.

## Code availability

Code and experiments are available at https://github.com/sriramlab/GPC.

## Data Availability

The 1000 Genomes datasets can be freely downloaded at https://www.internationalgenome.org/data. The UK Biobank dataset is available upon application at https://www.ukbiobank.ac.uk/.

## Footnotes

1 Note that [25] did not perform train/test splits so WGAN and RBM are actually trained on train+test.

## Notes

https://github.com/sriramlab/GPC

### Competing Interest Statement

The authors have declared no competing interest.

### Summary of Updates

Imputation Figures 5, 6, S5, and S6 now properly extend to 0.5 MAF; Author affiliations updated

