## Supplementary Information for "GPC: An expressive and tractable deep generative model for genetic variation data"

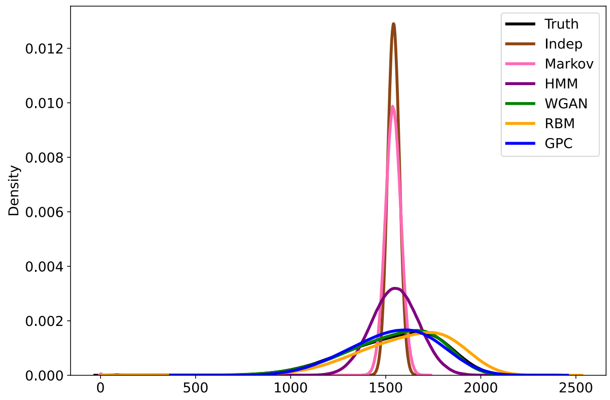


(a) Within each dataset (10K)

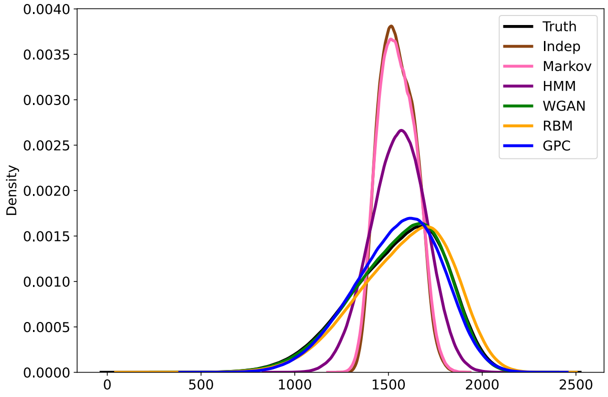


(b) Between datasets and test set (10K)

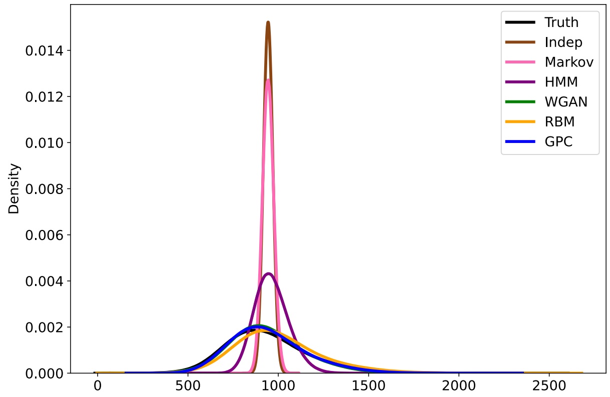


(c) Within each dataset (UKBB)

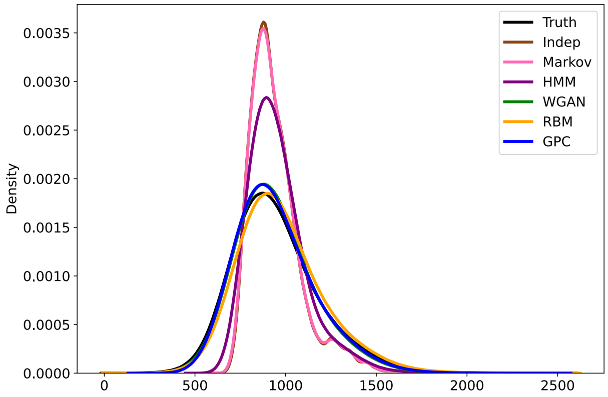


(d) Between datasets and test set (UKBB)

Supplementary Figure S1: **Distribution of haplotypic pairwise Euclidean distances.** (a–b) show results for the 1KG dataset; (c–d) for the UKBB dataset. (a,c) visualize distances *within* each dataset (i.e., intra-group variation). (b,d) show distances *between* the AG-generated samples and the test set (i.e., realism and mode collapse). Each panel compares across different generative models (see Table S1 for detailed metrics).

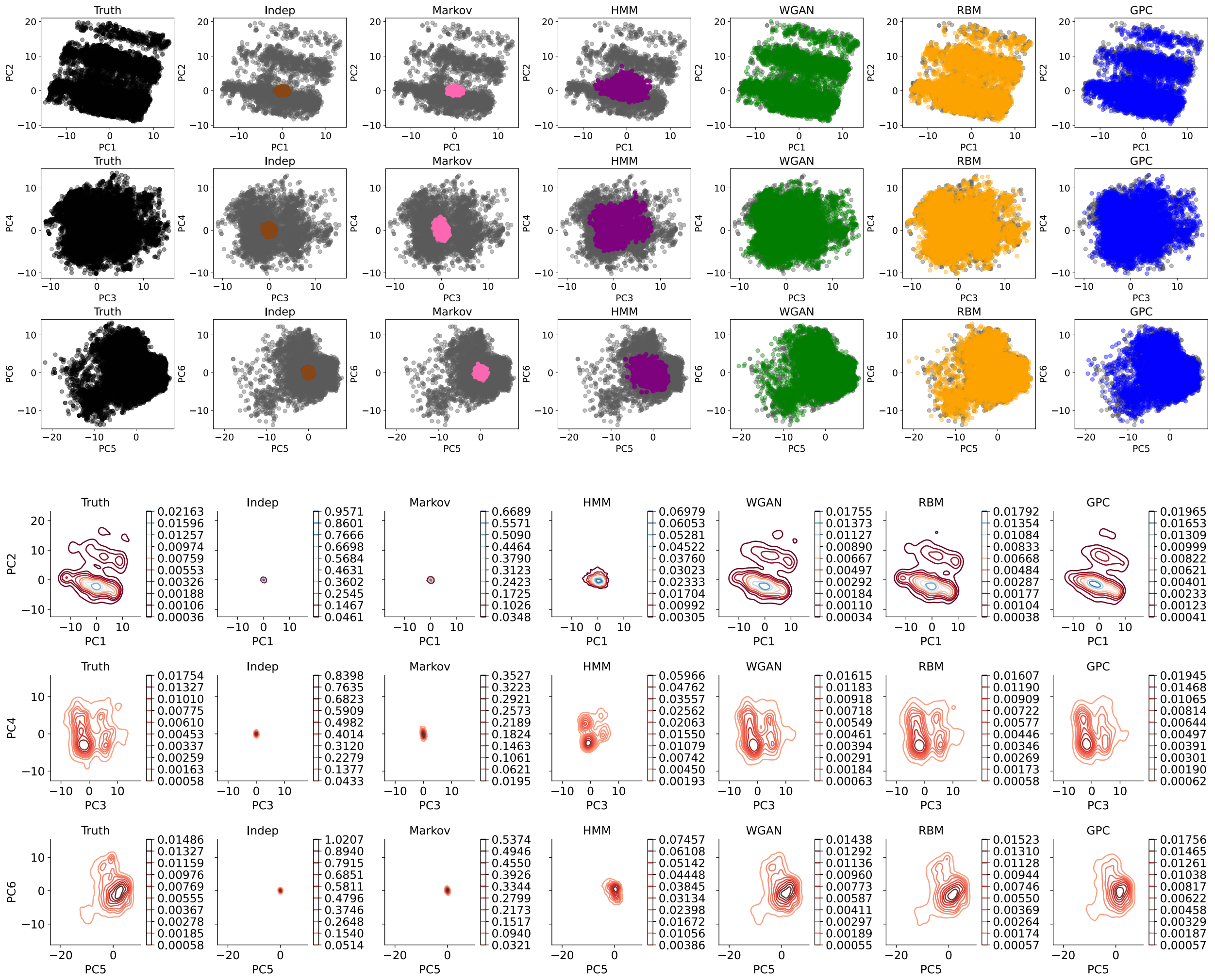


Supplementary Figure S2: **Principal components analysis for models trained on the UKBB dataset**. The top half shows a scatter plot of the top six principal components of the test set (gray) vs. AGs generated via Indep (brown), Markov (pink), HMM (purple), WGAN (green), RBM (orange), and GPC (blue). The left plot considers the train set as "perfectly" generated data (black). The bottom half of the figure shows a density map of the principal components for each dataset. All deep generative models are able to capture the population structure with good accuracy.

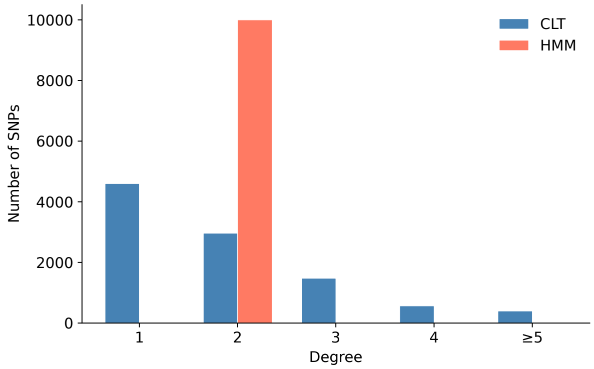


(a)

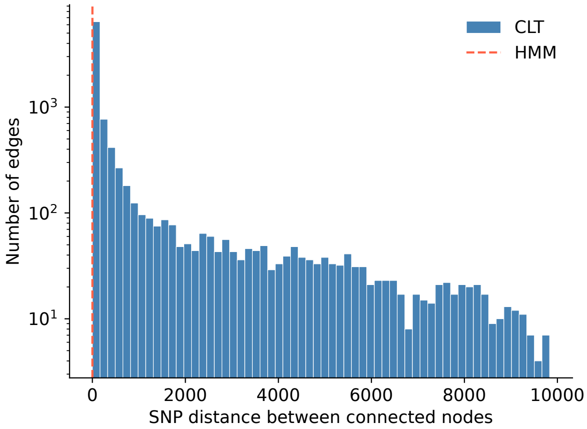


(b)

| **Property** | **CLT** | **HMM** |
|---|---|---|
| Total nodes (SNPs) | 10,000 | 10,000 |
| Leaf nodes (degree 1) | 4,599 | 2 |
| Degree-2 nodes | 2,962 | 9,998 |
| High-degree nodes (degree $\geq 5$) | 397 | 0 |
| Maximum node degree | 161 | 2 |
| Adjacent edges ($\|i-j\| = 1$) | 542 (5.4%) | 9,999 (100%) |
| Long-range edges ($\|i-j\| > 1{,}000$) | 1,834 (18.3%) | 0 (0%) |
| Maximum edge span ($\|i-j\|$) | 9,832 | 1 |

(c)

Supplementary Figure S3: **Comparison of the structures of Chow–Liu Tree (CLT) and a Hidden Markov Model (HMM) on the 1KG dataset. (a)** Degree distribution of CLT nodes versus HMM nodes. In an HMM chain over $N$ SNPs, exactly 2 nodes have degree 1 and all remaining $N-2$ nodes have degree 2. The CLT instead exhibits a highly heterogeneous degree distribution, with 4,599 leaf nodes, 2,962 degree-2 nodes, and 397 nodes with degree $\geq 5$, including one node (SNP 7579) with maximum degree 161. **(b)** Distribution of sequential edge spans ($|i-j|$ for each connected pair of SNPs), plotted on a log-scale $y$-axis. An HMM consists entirely of adjacent-SNP edges (dashed red line at distance 1), whereas the CLT spans distances up to 9,832 SNP positions, with 18.3% of edges connecting SNPs more than 1,000 positions apart. **(c)** Summary statistics comparing CLT and HMM topologies. Together, these panels demonstrate that the Chow–Liu algorithm learns a tree structure fundamentally distinct from a chain, directly linking SNPs with strong mutual information regardless of their genomic proximity and thereby encoding both local and long-range LD structure.

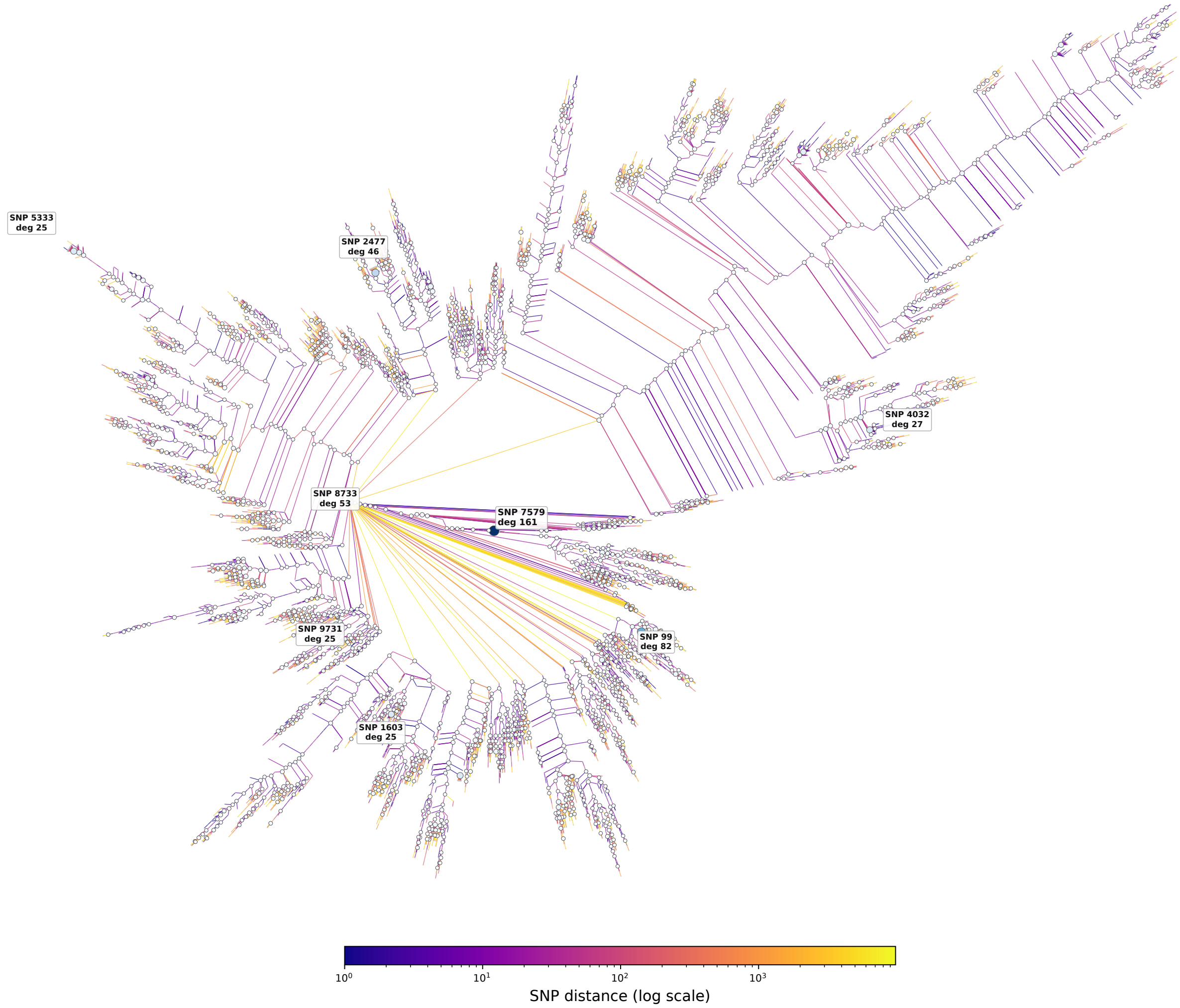


Supplementary Figure S4: **Radial visualization of the Chow–Liu Tree (CLT) learned from the 1KG dataset.** Each of the 10,000 nodes represents one SNP; edges connect pairs of SNPs whose latent variables are adjacent in the tree. Edge color encodes the sequential SNP distance $|i-j|$ on a log scale, ranging from deep purple (nearby SNPs, $|i-j| \approx 1$) to yellow (SNPs thousands of positions apart, $|i-j| \sim 10^3$). High-degree nodes (degree $\geq 3$) are shown as filled circles scaled by degree; leaf and degree-2 nodes appear as small gray circles. The top 8 highest-degree nodes are labeled with their SNP index and degree. In contrast to an HMM, which would appear as a single unbranched chain, the CLT exhibits a highly branching structure. For the highest-degree node SNP 7579 (degree 161), the edges leaving it are not visually apparent as individual lines because the vast majority of its 161 connected SNPs are themselves leaves with no further subtrees extending from them; they appear simply as the small gray circles immediately surrounding it. This contrasts with SNP 8733 (degree 53), whose connected nodes have large subtrees branching out from them, producing the visually prominent clusters that extend outward from that region of the tree. The long orange and yellow edges encode direct connections between SNPs thousands of genomic positions apart, capturing long-range LD that a chain topology cannot represent without propagating information through all intermediate variables.

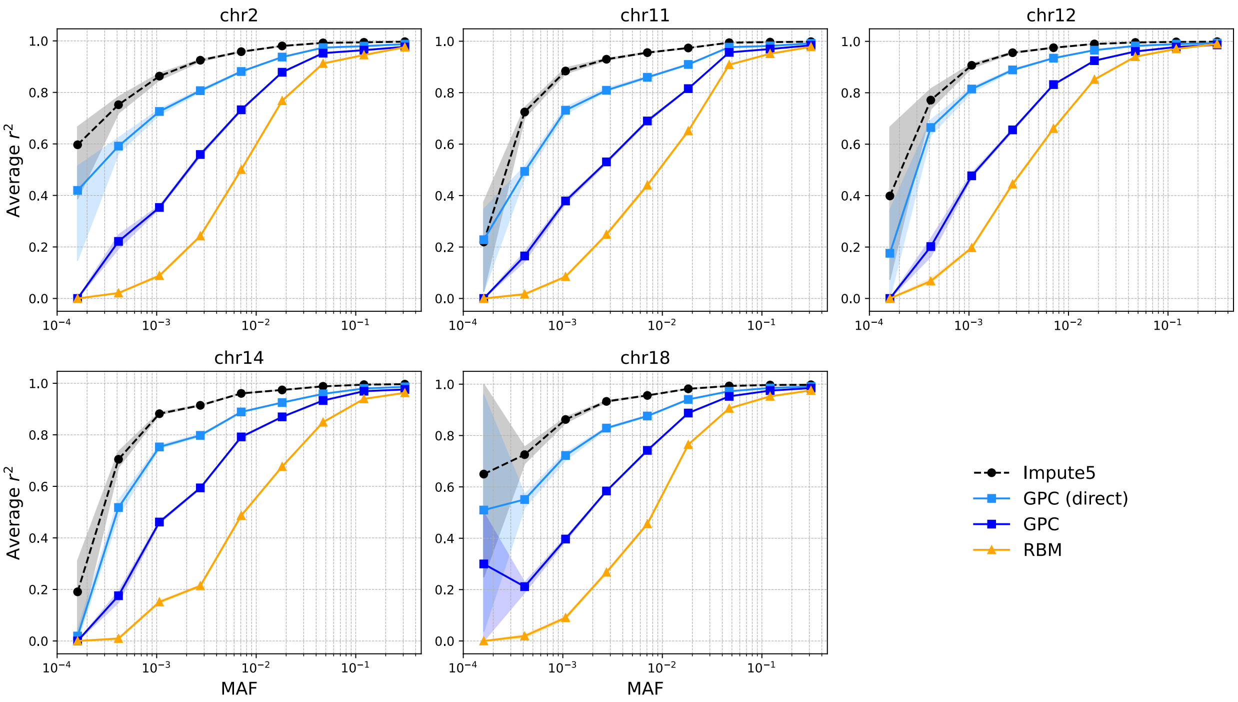


Supplementary Figure S5: **General imputation across additional UKBB regions.** The black and light blue lines denote imputation results using Impute5 with real reference genomes and direct imputation using GPC, respectively. The other lines show imputation results using Impute5 with AG reference panels. $r^2$ is shown as a function of MAF for each of the five additional chromosomes (chr2, 11, 12, 14, 18) separately (see Table S4 for detailed metrics).

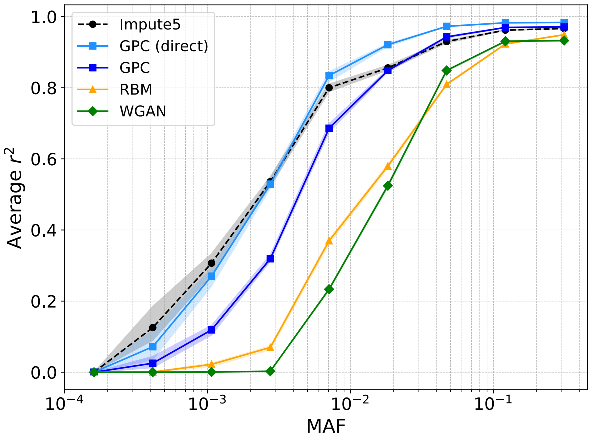


(a) Non-European target

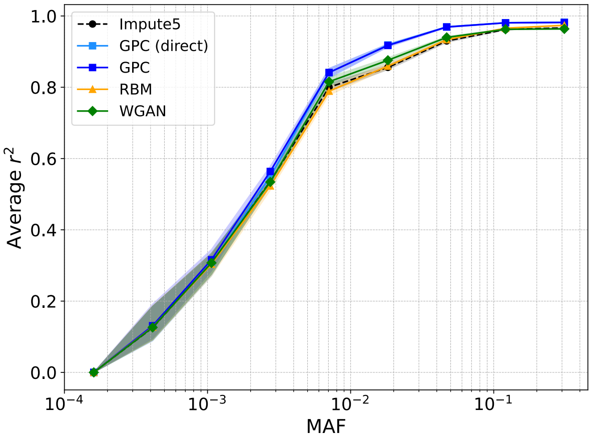


(b) Non-European target

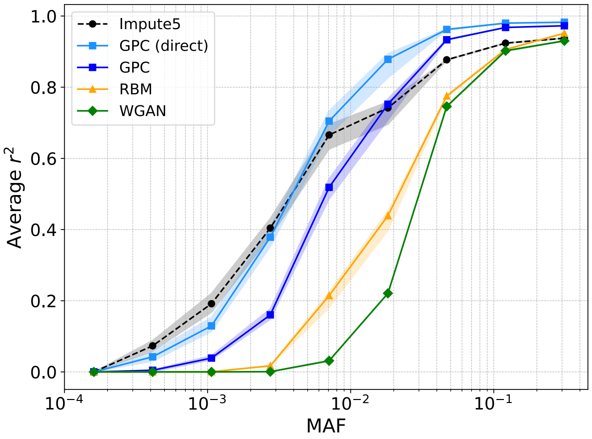


(c) African target

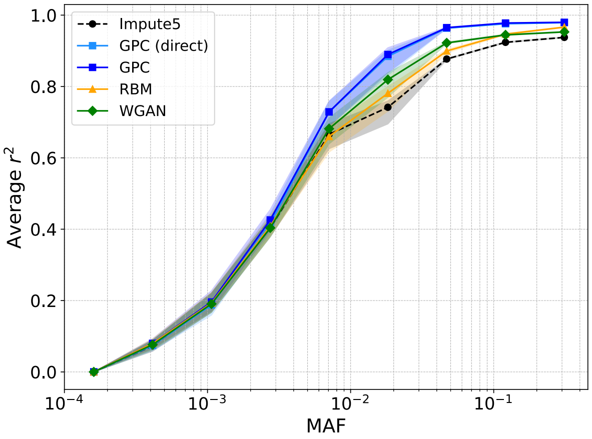


(d) African target

Supplementary Figure S6: **Population-specific imputation in the UKBB dataset.** On the left, the AG lines show results using population-specific AGs alone while the light blue line shows direct imputation with GPC trained on population-specific genomes (where the target genomes correspond to non-European populations in part a and African populations in part c). On the right, the AG lines show results using combined reference panels (real European data plus population-specific AGs), and the light blue line shows direct imputation with GPC trained on both European and population-specific genomes (where the target genomes correspond to non-European populations in part b and African populations in part d). The black line shows results using Impute5 with real European data as the reference panel (see Table S6 for detailed metrics).

| Dataset | | Truth | Indep | Markov | HMM | WGAN | RBM | GPC |
|---|---|---|---|---|---|---|---|---|
| **1KG** | Within | 10.66 | 180.23 | 172.70 | 105.88 | **14.73** | 61.99 | 23.59 |
| | Between | 5.34 | 128.24 | 125.61 | 86.25 | **6.35** | 31.50 | 18.73 |
| | PCA1-2 | 0.0026 | 0.2355 | 0.2085 | 0.0348 | 0.0028 | **0.0027** | 0.0039 |
| | PCA3-4 | 0.0022 | 0.1476 | 0.1148 | 0.0123 | **0.0023** | 0.0024 | 0.0028 |
| | PCA5-6 | 0.0023 | 0.1011 | 0.0822 | 0.0042 | 0.0024 | **0.0023** | 0.0029 |
| **UKBB** | Within | 3.57 | 168.88 | 164.55 | 117.91 | 25.16 | 49.52 | **20.86** |
| | Between | 1.81 | 80.20 | 79.36 | 64.25 | 13.08 | 25.81 | **11.88** |
| | PCA1-2 | 0.0019 | 0.0909 | 0.0846 | 0.0072 | 0.0022 | **0.0021** | 0.0024 |
| | PCA3-4 | 0.0019 | 0.0656 | 0.0634 | 0.0244 | 0.0022 | **0.0022** | 0.0026 |
| | PCA5-6 | 0.0019 | 0.0634 | 0.0542 | 0.0117 | 0.0021 | **0.0020** | 0.0023 |

Supplementary Table S1: **Distance metrics between real and generated data. Within, Between** (Figure S1): Within denotes the Wasserstein distance between the distributions of pairwise haplotypic distances computed within a single dataset (e.g., generated–generated) and the corresponding distribution computed within the real test set (real–real). Between denotes the Wasserstein distance between the distribution of pairwise haplotypic distances computed between generated and real test individuals (generated–real) and the distribution computed within the real test set (real–real). In both cases, the real test set serves as the reference distribution. **PCA1-2, PCA3-4, PCA5-6**: Wasserstein 2D distances between the PCA representations of real (test set) versus generated individuals. Truth represents the training set. For each PCA distance, one PCA is fit on the method AGs and test data. Bolded values indicate the best among all compared models.

| **Dataset** | **Metric** | Truth | Indep | Markov | HMM | WGAN | RBM | GPC |
|---|---|---|---|---|---|---|---|---|
| **1KG** | MAE | 0.0008 | 0.1676 | 0.0708 | 0.0153 | 0.0237 | 0.0182 | **0.0053** |
| | RMSE | 0.0011 | 0.2101 | 0.1031 | 0.0240 | 0.0361 | 0.0307 | **0.0056** |
| **UKBB** | MAE | 0.0003 | 0.1111 | 0.0481 | 0.0153 | 0.0186 | 0.0295 | **0.0047** |
| | RMSE | 0.0004 | 0.1336 | 0.0649 | 0.0262 | 0.0283 | 0.0414 | **0.0052** |

Supplementary Table S2: **LD decay error metrics for Figure 3**. Mean absolute error (MAE) and root mean squared error (RMSE) are computed as differences between the mean values of each bin on the LD decay curves across genomic distance. Lower values indicate better preservation of LD structure; best-performing values are bolded.

**1KG**

| **Method** | **Mean $r^2$ [95% CI]** |
|---|---|
| **All SNPs** | |
| Impute5 | 0.649 [0.643, 0.655] |
| RBM (full) | 0.510 [0.506, 0.514] |
| WGAN (full) | 0.428 [0.426, 0.430] |
| RBM | 0.494 [0.491, 0.496] |
| WGAN | 0.445 [0.443, 0.447] |
| GPC | 0.540 [0.536, 0.544] |
| **GPC (direct)** | **0.593 [0.589, 0.598]** |
| **Low-freq** (MAF $< 1\%$) | |
| Impute5 | 0.298 [0.286, 0.310] |
| RBM (full) | 0.095 [0.088, 0.103] |
| WGAN (full) | 0.004 [0.004, 0.005] |
| RBM | 0.066 [0.060, 0.072] |
| WGAN | 0.009 [0.009, 0.010] |
| GPC | 0.132 [0.123, 0.140] |
| **GPC (direct)** | **0.218 [0.208, 0.228]** |
| **Rare** (MAF $< 0.1\%$) | |
| Impute5 | 0.089 [0.078, 0.099] |
| RBM (full) | 0.012 [0.007, 0.017] |
| WGAN (full) | 0.000 [0.000, 0.000] |
| RBM | 0.007 [0.005, 0.009] |
| WGAN | 0.001 [0.000, 0.001] |
| GPC | 0.012 [0.008, 0.017] |
| **GPC (direct)** | **0.053 [0.044, 0.061]** |

**UKBB**

| **Method** | **Mean $r^2$ [95% CI]** |
|---|---|
| **All SNPs** | |
| Impute5 | 0.957 [0.954, 0.960] |
| RBM | 0.669 [0.667, 0.670] |
| WGAN | 0.523 [0.522, 0.524] |
| GPC | 0.750 [0.748, 0.751] |
| **GPC (direct)** | **0.906 [0.903, 0.909]** |
| **Low-freq** (MAF $< 1\%$) | |
| Impute5 | 0.914 [0.907, 0.922] |
| RBM | 0.381 [0.377, 0.384] |
| WGAN | 0.087 [0.086, 0.088] |
| GPC | 0.525 [0.522, 0.528] |
| **GPC (direct)** | **0.828 [0.822, 0.835]** |
| **Rare** (MAF $< 0.1\%$) | |
| Impute5 | 0.818 [0.796, 0.839] |
| RBM | 0.118 [0.112, 0.123] |
| WGAN | 0.000 [0.000, 0.000] |
| GPC | 0.263 [0.252, 0.274] |
| **GPC (direct)** | **0.709 [0.687, 0.730]** |

Supplementary Table S3: **Summarized metrics for Figure 5.** Mean imputation performance across 10 bootstrapped replicates for all, low-frequency (MAF $< 1\%$), and rare (MAF $< 0.1\%$) SNPs. Best generative model is bolded.

### Chromosome 2

| Method | Mean $r^2$ [95% CI] |
|---|---|
| **All SNPs** | |
| Impute5 | 0.949 [0.945, 0.952] |
| RBM | 0.605 [0.603, 0.606] |
| GPC | 0.746 [0.744, 0.748] |
| **GPC (direct)** | **0.884 [0.882, 0.887]** |
| **Low-freq** (MAF $<$ 1%) | |
| Impute5 | 0.894 [0.886, 0.902] |
| RBM | 0.222 [0.219, 0.225] |
| GPC | 0.491 [0.486, 0.496] |
| **GPC (direct)** | **0.774 [0.768, 0.780]** |
| **Rare** (MAF $<$ 0.1%) | |
| Impute5 | 0.805 [0.782, 0.827] |
| RBM | 0.043 [0.037, 0.048] |
| GPC | 0.269 [0.253, 0.286] |
| **GPC (direct)** | **0.654 [0.635, 0.673]** |

### Chromosome 11

| Method | Mean $r^2$ [95% CI] |
|---|---|
| **All SNPs** | |
| Impute5 | 0.959 [0.956, 0.962] |
| RBM | 0.640 [0.639, 0.642] |
| GPC | 0.770 [0.768, 0.772] |
| **GPC (direct)** | **0.895 [0.891, 0.898]** |
| **Low-freq** (MAF $<$ 1%) | |
| Impute5 | 0.903 [0.896, 0.911] |
| RBM | 0.214 [0.211, 0.218] |
| GPC | 0.484 [0.478, 0.490] |
| **GPC (direct)** | **0.769 [0.760, 0.778]** |
| **Rare** (MAF $<$ 0.1%) | |
| Impute5 | 0.801 [0.778, 0.824] |
| RBM | 0.033 [0.030, 0.035] |
| GPC | 0.272 [0.257, 0.287] |
| **GPC (direct)** | **0.621 [0.598, 0.644]** |

### Chromosome 12

| Method | Mean $r^2$ [95% CI] |
|---|---|
| **All SNPs** | |
| Impute5 | 0.970 [0.968, 0.973] |
| RBM | 0.717 [0.715, 0.718] |
| GPC | 0.823 [0.821, 0.824] |
| **GPC (direct)** | **0.934 [0.931, 0.937]** |
| **Low-freq** (MAF $<$ 1%) | |
| Impute5 | 0.937 [0.932, 0.942] |
| RBM | 0.423 [0.419, 0.426] |
| GPC | 0.636 [0.632, 0.639] |
| **GPC (direct)** | **0.869 [0.863, 0.876]** |
| **Rare** (MAF $<$ 0.1%) | |
| Impute5 | 0.823 [0.795, 0.851] |
| RBM | 0.115 [0.105, 0.126] |
| GPC | 0.330 [0.314, 0.347] |
| **GPC (direct)** | **0.723 [0.696, 0.750]** |

### Chromosome 14

| Method | Mean $r^2$ [95% CI] |
|---|---|
| **All SNPs** | |
| Impute5 | 0.946 [0.943, 0.948] |
| RBM | 0.569 [0.567, 0.571] |
| GPC | 0.761 [0.759, 0.763] |
| **GPC (direct)** | **0.878 [0.874, 0.881]** |
| **Low-freq** (MAF $<$ 1%) | |
| Impute5 | 0.897 [0.891, 0.903] |
| RBM | 0.243 [0.240, 0.247] |
| GPC | 0.561 [0.557, 0.566] |
| **GPC (direct)** | **0.781 [0.774, 0.788]** |
| **Rare** (MAF $<$ 0.1%) | |
| Impute5 | 0.790 [0.765, 0.814] |
| RBM | 0.053 [0.047, 0.058] |
| GPC | 0.293 [0.278, 0.308] |
| **GPC (direct)** | **0.627 [0.603, 0.651]** |

### Chromosome 18

| Method | Mean $r^2$ [95% CI] |
|---|---|
| **All SNPs** | |
| Impute5 | 0.950 [0.947, 0.953] |
| RBM | 0.598 [0.596, 0.599] |
| GPC | 0.759 [0.757, 0.762] |
| **GPC (direct)** | **0.886 [0.883, 0.890]** |
| **Low-freq** (MAF $<$ 1%) | |
| Impute5 | 0.897 [0.890, 0.904] |
| RBM | 0.222 [0.218, 0.225] |
| GPC | 0.523 [0.517, 0.528] |
| **GPC (direct)** | **0.779 [0.773, 0.786]** |
| **Rare** (MAF $<$ 0.1%) | |
| Impute5 | 0.786 [0.759, 0.813] |
| RBM | 0.037 [0.029, 0.044] |
| GPC | 0.293 [0.276, 0.310] |
| **GPC (direct)** | **0.631 [0.605, 0.657]** |

Supplementary Table S4: **Summarized metrics for Supplementary Figure S5 (additional UKBB regions).** Mean imputation performance across 10 bootstrapped replicates for all, low-frequency (MAF $<$ 1%), and rare (MAF $<$ 0.1%) SNPs on five additional LD blocks in the UKBB dataset. Best generative model is bolded.

**Non-European**

| Method | Mean $r^2$ [95% CI] |
|---|---|
| **All SNPs** | |
| Impute5 (EUR) | 0.485 [0.482, 0.488] |
| RBM | 0.476 [0.474, 0.478] |
| WGAN | 0.422 [0.421, 0.424] |
| GPC | 0.522 [0.520, 0.524] |
| GPC (direct)† | 0.563 [0.561, 0.564]† |
| RBM (combined) | 0.528 [0.524, 0.532] |
| WGAN (combined) | 0.507 [0.504, 0.510] |
| GPC (combined) | 0.560 [0.555, 0.562] |
| **GPC (direct combined)** | **0.570 [0.567, 0.573]** |
| **Low-freq** (MAF < 1%) | |
| Impute5 (EUR) | 0.106 [0.101, 0.112] |
| RBM | 0.042 [0.039, 0.046] |
| WGAN | 0.003 [0.002, 0.004] |
| GPC | 0.098 [0.095, 0.101] |
| GPC (direct)† | 0.158 [0.156, 0.161]† |
| RBM (combined) | 0.123 [0.116, 0.129] |
| WGAN (combined) | 0.111 [0.105, 0.117] |
| GPC (combined) | 0.160 [0.155, 0.165] |
| **GPC (direct combined)** | **0.174 [0.169, 0.178]** |
| **Rare** (MAF < 0.1%) | |
| Impute5 (EUR) | 0.021 [0.016, 0.025] |
| RBM | 0.003 [0.001, 0.004] |
| WGAN | 0.000 [0.000, 0.001] |
| GPC | 0.004 [0.002, 0.006] |
| GPC (direct)† | 0.021 [0.016, 0.026]† |
| RBM (combined) | 0.024 [0.019, 0.029] |
| WGAN (combined) | 0.021 [0.016, 0.025] |
| GPC (combined) | 0.026 [0.021, 0.030] |
| **GPC (direct combined)** | **0.031 [0.025, 0.036]** |

**African**

| Method | Mean $r^2$ [95% CI] |
|---|---|
| **All SNPs** | |
| Impute5 (EUR) | 0.368 [0.357, 0.379] |
| RBM | 0.429 [0.424, 0.434] |
| WGAN | 0.381 [0.379, 0.382] |
| GPC | 0.442 [0.438, 0.446] |
| GPC (direct)† | 0.462 [0.456, 0.468]† |
| RBM (combined) | 0.446 [0.435, 0.456] |
| WGAN (combined) | 0.423 [0.412, 0.435] |
| GPC (combined) | 0.464 [0.454, 0.474] |
| **GPC (direct combined)** | **0.471 [0.461, 0.480]** |
| **Low-freq** (MAF < 1%) | |
| Impute5 (EUR) | 0.019 [0.013, 0.025] |
| RBM | 0.028 [0.025, 0.030] |
| WGAN | 0.009 [0.008, 0.009] |
| GPC | 0.027 [0.025, 0.028] |
| GPC (direct)† | 0.043 [0.040, 0.046]† |
| RBM (combined) | 0.040 [0.033, 0.047] |
| WGAN (combined) | 0.028 [0.022, 0.035] |
| GPC (combined) | 0.043 [0.037, 0.050] |
| **GPC (direct combined)** | **0.053 [0.046, 0.059]** |
| **Rare** (MAF < 0.1%) | |
| Impute5 (EUR) | 0.003 [0.001, 0.005] |
| RBM | 0.002 [0.001, 0.003] |
| WGAN | 0.000 [0.000, 0.001] |
| GPC | 0.001 [0.001, 0.001] |
| GPC (direct)† | 0.006 [0.005, 0.008]† |
| RBM (combined) | 0.004 [0.002, 0.006] |
| WGAN (combined) | 0.003 [0.001, 0.005] |
| GPC (combined) | 0.004 [0.002, 0.005] |
| **GPC (direct combined)** | **0.007 [0.005, 0.009]** |

Supplementary Table S5: **Summarized metrics for Figure 6 (1KG).** Mean imputation performance across 10 bootstrapped replicates for all, low-frequency (MAF < 1%), and rare (MAF < 0.1%) SNPs. Best generative model using combined data is bolded; best using population-specific data only is marked with †.

**Non-European**

| **Method** | **Mean $r^2$ [95% CI]** |
|---|---|
| **All SNPs** | |
| Impute5 (EUR) | 0.733 [0.725, 0.742] |
| RBM | 0.495 [0.493, 0.497] |
| WGAN | 0.461 [0.460, 0.463] |
| GPC | 0.656 [0.651, 0.661] |
| GPC (direct)† | 0.749 [0.742, 0.757]† |
| RBM (combined) | 0.732 [0.723, 0.740] |
| WGAN (combined) | 0.739 [0.731, 0.748] |
| **GPC (combined)** | **0.763 [0.755, 0.771]** |
| GPC (direct combined) | 0.759 [0.752, 0.767] |
| **Low-freq** (MAF $<$ 1%) | |
| Impute5 (EUR) | 0.506 [0.491, 0.522] |
| RBM | 0.128 [0.123, 0.132] |
| WGAN | 0.059 [0.058, 0.061] |
| GPC | 0.333 [0.323, 0.343] |
| GPC (direct)† | 0.498 [0.483, 0.513]† |
| RBM (combined) | 0.498 [0.482, 0.514] |
| WGAN (combined) | 0.510 [0.494, 0.526] |
| **GPC (combined)** | **0.530 [0.515, 0.546]** |
| GPC (direct combined) | 0.522 [0.507, 0.537] |
| **Rare** (MAF $<$ 0.1%) | |
| Impute5 (EUR) | 0.211 [0.174, 0.248] |
| RBM | 0.004 [0.003, 0.006] |
| WGAN | 0.000 [0.000, 0.000] |
| GPC | 0.054 [0.034, 0.073] |
| GPC (direct)† | 0.159 [0.127, 0.190]† |
| RBM (combined) | 0.210 [0.171, 0.248] |
| WGAN (combined) | 0.209 [0.173, 0.245] |
| **GPC (combined)** | **0.217 [0.180, 0.255]** |
| GPC (direct combined) | 0.210 [0.175, 0.246] |

**African**

| **Method** | **Mean $r^2$ [95% CI]** |
|---|---|
| **All SNPs** | |
| Impute5 (EUR) | 0.645 [0.627, 0.663] |
| RBM | 0.437 [0.428, 0.446] |
| WGAN | 0.375 [0.374, 0.376] |
| GPC | 0.582 [0.571, 0.593] |
| GPC (direct)† | 0.679 [0.663, 0.694]† |
| RBM (combined) | 0.659 [0.641, 0.678] |
| WGAN (combined) | 0.668 [0.650, 0.686] |
| **GPC (combined)** | **0.702 [0.684, 0.720]** |
| GPC (direct combined) | 0.698 [0.679, 0.716] |
| **Low-freq** (MAF $<$ 1%) | |
| Impute5 (EUR) | 0.384 [0.354, 0.414] |
| RBM | 0.061 [0.052, 0.070] |
| WGAN | 0.009 [0.008, 0.009] |
| GPC | 0.207 [0.191, 0.224] |
| GPC (direct)† | 0.364 [0.337, 0.390]† |
| RBM (combined) | 0.383 [0.352, 0.415] |
| WGAN (combined) | 0.388 [0.358, 0.418] |
| **GPC (combined)** | **0.410 [0.380, 0.441]** |
| GPC (direct combined) | 0.404 [0.373, 0.434] |
| **Rare** (MAF $<$ 0.1%) | |
| Impute5 (EUR) | 0.115 [0.091, 0.139] |
| RBM | 0.001 [0.000, 0.001] |
| WGAN | 0.000 [0.000, 0.000] |
| GPC | 0.015 [0.008, 0.023] |
| GPC (direct)† | 0.064 [0.051, 0.078]† |
| RBM (combined) | 0.117 [0.094, 0.139] |
| WGAN (combined) | 0.115 [0.093, 0.138] |
| **GPC (combined)** | **0.118 [0.095, 0.141]** |
| GPC (direct combined) | 0.111 [0.087, 0.135] |

Supplementary Table S6: **Summarized metrics for Figure S6 (UKBB).** Mean imputation performance across 10 bootstrapped replicates for all, low-frequency (MAF $<$ 1%), and rare (MAF $<$ 0.1%) SNPs. Best generative model using combined data is bolded; best using population-specific data only is marked with †.

| **Method** | **Mean $r^2$ [95% CI]** |
| --- | --- |
| **All SNPs** | |
| Impute5 | 0.749 [0.745, 0.753] |
| RBM | 0.574 [0.570, 0.578] |
| WGAN | 0.474 [0.472, 0.476] |
| GPC | 0.515 [0.511, 0.518] |
| **GPC (direct)** | **0.594 [0.589, 0.598]** |
| **Low-freq** (MAF $< 1\%$) | |
| Impute5 | 0.626 [0.618, 0.633] |
| RBM | 0.301 [0.290, 0.312] |
| WGAN | 0.120 [0.116, 0.123] |
| GPC | 0.215 [0.209, 0.222] |
| **GPC (direct)** | **0.344 [0.335, 0.352]** |

Supplementary Table S7: **Summarized metrics for Figure 7 (high-coverage 1KG).** Mean imputation performance across 10 bootstrapped replicates for all and low-frequency (MAF $< 1\%$) SNPs. Best generative model is bolded.

**Non-European**

| Method | Mean $r^2$ [95% CI] |
|---|---|
| **All SNPs** | |
| Impute5 (EUR) | 0.338 [0.335, 0.342] |
| RBM | 0.565 [0.562, 0.568] |
| WGAN | 0.462 [0.459, 0.466] |
| GPC | 0.529 [0.527, 0.531] |
| GPC (direct)† | 0.593 [0.590, 0.596]† |
| RBM (combined) | 0.584 [0.580, 0.588] |
| WGAN (combined) | 0.494 [0.489, 0.499] |
| GPC (combined) | 0.548 [0.545, 0.552] |
| **GPC (direct combined)** | **0.584 [0.581, 0.586]** |
| **Low-freq** (MAF < 1%) | |
| Impute5 (EUR) | 0.046 [0.042, 0.049] |
| RBM | 0.296 [0.287, 0.304] |
| WGAN | 0.114 [0.112, 0.116] |
| GPC | 0.237 [0.234, 0.240] |
| GPC (direct)† | 0.339 [0.335, 0.344]† |
| RBM (combined) | 0.329 [0.321, 0.337] |
| WGAN (combined) | 0.162 [0.155, 0.169] |
| GPC (combined) | 0.266 [0.256, 0.276] |
| **GPC (direct combined)** | **0.331 [0.323, 0.338]** |

**African**

| Method | Mean $r^2$ [95% CI] |
|---|---|
| **All SNPs** | |
| Impute5 (EUR) | 0.268 [0.264, 0.273] |
| RBM | 0.552 [0.543, 0.562] |
| WGAN | 0.367 [0.363, 0.371] |
| GPC | 0.546 [0.539, 0.552] |
| GPC (direct)† | 0.583 [0.574, 0.591]† |
| **RBM (combined)** | **0.563 [0.553, 0.574]** |
| WGAN (combined) | 0.407 [0.398, 0.416] |
| GPC (combined) | 0.556 [0.548, 0.564] |
| GPC (direct combined) | 0.562 [0.552, 0.571] |
| **Low-freq** (MAF < 1%) | |
| Impute5 (EUR) | 0.022 [0.017, 0.027] |
| RBM | 0.339 [0.320, 0.359] |
| WGAN | 0.082 [0.078, 0.086] |
| GPC | 0.317 [0.299, 0.334] |
| GPC (direct)† | 0.376 [0.355, 0.396]† |
| **RBM (combined)** | **0.350 [0.330, 0.371]** |
| WGAN (combined) | 0.101 [0.093, 0.110] |
| GPC (combined) | 0.325 [0.307, 0.344] |
| GPC (direct combined) | 0.342 [0.320, 0.363] |

Supplementary Table S8: **Summarized metrics for Figure 8 (high-coverage 1KG).** Mean imputation performance across 10 bootstrapped replicates for all and low-frequency (MAF < 1%) SNPs. Best generative model using combined data is bolded; best using population-specific data only is marked with †. In this experiment, GPC (direct)† performs better than models using combined data.
